# HINT1 Inhibitors as Selective Modulators of MOR-NMDAR Cross Regulation and Non-Opioid Analgesia

**DOI:** 10.1101/2024.07.15.603568

**Authors:** Maxwell Dillenburg, Cristina D. Peterson, Rafal Dolat, Kostana Ligori, Kelley F Kitto, George L Wilcox, Carolyn A. Fairbanks, Carston R. Wagner

## Abstract

The Human Histidine Triad Nucleotide Binding Protein 1 (HINT1) has recently become a protein of interest due to its involvement in several CNS processes, including neuroplasticity and the development of several neuropsychiatric disorders. Crucially, HINT1 behaves as a mediator for the cross-regulation of the mu opioid receptor (MOR) and *N*-methyl-D-aspartate receptor (NMDAR). Active site inhibition of HINT1 using small molecule inhibitors has been demonstrated to have a significant impact on this cross-regulatory relationship *in vivo*. Herein, we describe the development of a series of ethenoadenosine HINT1 inhibitors to further evaluate the effect of HINT1 inhibition on morphine’s blockade of NMDA-evoked behaviors, the development of acute endomorphin-2 tolerance and analgesia. X-ray crystallographic analysis and HINT1 binding experiments demonstrate that modifications to the inhibitor nucleobase greatly impact the inhibitor binding interactions with HINT1. Our results reveal a complex structural-activity relationship for HINT1 inhibitors in which minor modifications to the ethenoadenosine scaffold resulted in dramatic changes to their activity in these assays modeling MOR-NMDAR interaction. Specifically, we observed the ability of HINT1 inhibitors to selectively affect individual pathways of MOR-NMDAR crosstalk. Furthermore, we observed that a carbamate ethenoadenosine inhibitor of HINT1 can induce analgesia, while not affecting opioid tolerance. Additionally, although past studies have indicated that that loss of HINT1 expression can result in the downregulation of p53, we have shown that inhibition of HINT1 has no effect on either the expression of HINT1 or p53. These studies highlight the critical role of HINT1 in MOR-NMDAR crosstalk and demonstrate the intriguing potential of using HINT1 active-site inhibitors as tools to probe its role in these biochemical pathways and its potential as a novel pain target.

## Introduction

HINT1 has recently garnered interest due to its involvement in several CNS processes and its association with the development of multiple neuropsychiatric disorders.^1–2^ Additionally, targeting HINT1 using small molecule HINT1 inhibitors has demonstrated its potential as a target for pain therapy.^3–4^ The histidine triad nucleotide binding protein 1 (HINT1) is a member of the ubiquitously expressed and ancient superfamily of histidine triad (HIT) proteins, which are characterized by their conserved catalytic sequence His-X-His-X-His-X-X, where X is a hydrophobic residue. HINT1 is a 14 kD homodimer that possesses phosphoramidase and acyl-AMP hydrolase activity, with a preference for purine over pyrimidine nucleoside substrates.^5–6^ This catalytic activity is crucial for the activation of several clinically relevant nucleoside phosphoramidate prodrugs, but the endogenous function and substrate of HINT1 are not well understood.^5, 7^ While the exact role of HINT1’s enzymatic activity still remains a mystery, HINT1 is observed to participate in wide array of biological processes such as tumor suppression, mast cell activation, and apoptosis.^8–11^ HINT1 participates in these processes via interaction with transcription factor complexes such as pontin and reptin, LEC/TCF, and MITF/USF2.^8–11^ The involvement of HINT1 in such a range of processes makes it an exciting target for further evaluation.

HINT1 has widespread expression in the CNS, with the high levels observed in cerebral cortex, periaqueductal gray area, and nucleus accumbens, which are responsible for motor and sensory functions, modulation of pain, and addiction properties respectively.^12^ Alterations to HINT1 expression and HINT1 mutations have been associated with the development of several neuropsychiatric disorders including schizophrenia, addiction, and inherited peripheral neuropathies.^1, 13–16^ HINT1^-/-^ mice display several behavioral changes compared to their wildtype counterparts, including decreased nicotine dependence, hypersensitivity to amphetamines, and increased anxiety and depression-like behaviors, demonstrating a crucial role for HINT1 in several CNS processes.^13–14, 17–18^ Further, HINT1 regulates the signaling of several CNS receptors such as the mu opioid receptor (MOR), cannabinoid receptors, transient receptor potential cation channels, and sigma receptors, making it an extremely interesting target for pharmacological interrogation.^19–22^ In the case of MOR, HINT1 is critical in mediating the cross-regulation observed between MOR and the N-methyl-D-aspartate receptor (NMDAR).^4, 21^ Following opioid activation of MOR and its subsequent analgesic pathway, HINT1 mediates the formation of several protein assemblies which lead to the activation of NMDAR.^2, 23^ The resulting signaling cascade ultimately leads to phosphorylation of MOR and downregulation of its signaling.^21, 23^

Previously, our lab has developed small molecule HINT1 inhibitors via replacement of hydrolysable phosphoramidate and acyl-monophosphate groups of compounds **1** and **2** with non-hydrolysable bioisosteres such as the carbamates and acyl-sulfamates, as shown by **TrpGc** and **3**, respectively (**Figure 1**).^3–4^ We have demonstrated that active-site inhibition of HINT1 using these nucleoside-based inhibitors has a striking effect on several pain pathways *in vivo*. Treatment of mice with morphine and our HINT1 inhibitor **TrpGc** results in an increased analgesic response and decreased development of morphine tolerance.^3–4^ Importantly, these effects were not observed in HINT1^-/-^ mice following treatment with **TrpGc**.^3^ Assessment of a series of HINT1 inhibitors in two forms of MOR-NMDAR crosstalk, namely MOR inhibition of NMDAR activation and endomorphin-2 tolerance, demonstrated the first SAR of HINT1 inhibition and continued to establish the stark pharmacological effect of HINT1 inhibition in these processes.^4^ Notably, the tryptamine side chain of our HINT1 inhibitors does not contribute significantly to HINT1 binding, but is crucial to the *in vivo* activity of HINT1 inhibitors.^4^ Additionally, both carbamate and acyl-sulfamate based HINT1 inhibitors were capable of blocking MOR inhibition of NMDAR evoked thermal hypersensitivity, but only the carbamate based **TrpGc** blocked the development of endomorphin-2 tolerance.^4^ Interestingly, we found that the binding affinity of HINT1 inhibitors did not correlate to their activity *in vivo*. Importantly, we observed that active-site inhibition of HINT1 with small molecules can have selective effects across these assays, though the mechanism behind these effects is not clear.

**Figure 1.**
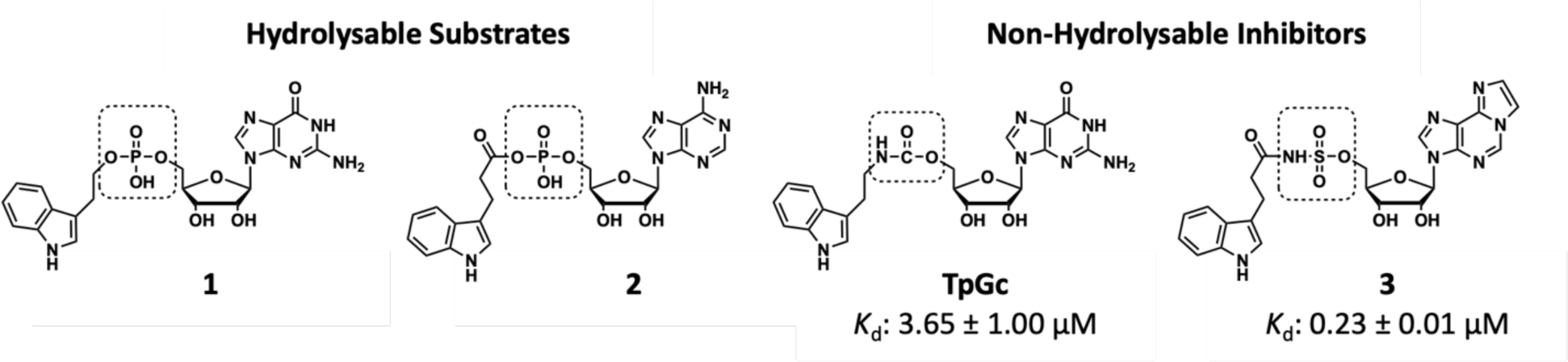
HINT1 Substrates and Previously Developed HINT1 Inhibitors. Notable HINT1 substrates and inhibitors. Dissociation constants for **TrpGc** and **3** binding to HINT1 were previously reported.^3, 23^

With the intriguing results from our previous SAR studies, specifically the ability of certain inhibitors to selectively impact individual pathways in MOR-NMDAR crosstalk, we sought to expand our arsenal of HINT1 inhibitors and further probe the pharmacological effects of HINT1 inhibition. Due to the high binding affinity and selective activity of our previously reported ethenoadenosine-based inhibitor, compound **3**, we have designed and synthesized a series of substituted ethenoadenosine-based inhibitors.^4, 23^ Importantly, due to the differential activities of the carbamate and acyl-sulfamate backbones, we have synthesized both analogues for each inhibitor. We have evaluated the binding of each of these inhibitors via their inhibition of HINT1 hydrolysis using our previously reported continuous fluorescence assay.^24^ Additionally, we analyzed the binding interactions of these inhibitors via X-ray crystallographic analysis. Finally, we evaluated each of these inhibitors for their ability to block MOR inhibition of NMDAR activation, prevent endomorphin-2 tolerance, and produce an analgesic response following spinal administration. Lastly, since HINT1 expression, but not enzymatic activity, has been shown to correlate with p53 expression, the effect of HINT1 inhibition on p53 expression was determined.^25^ Taken together, these results highlight the intriguing role of HINT1 in MOR-NMDAR crosstalk and the pharmacological opportunities provided by small molecule inhibition of HINT1.

## Results

### Modifications to the Ethenoadenosine Base Alter Inhibitor Binding to HINT1

We began by examining the impact of substitution to the ethenoadenosine base for our nucleoside carbamate inhibitors by evaluating their inhibition of HINT1 enzymatic activity. Due to the hydrogen bond observed between the 2-amino group of the guanosine base of HINT1 inhibitors and the backbone carbonyl of His42 of HINT1, we hypothesized that addition of an amine to the 2-position of the ethenoadenosine base could result in a similar hydrogen bond and improved inhibitor binding affinity.^4^ To probe the SAR of this position, we designed a series of three ethenoadenosine-based carbamate inhibitors (**compounds 8-10)** with varying degrees of substitution at the 2-position. Synthesis of **8** and **9** proceeded via the common intermediate **5**. We proceeded beginning with the commercially available 2-chloroadenosine. First, acetonide protection of the 2’, 3’ – OH was achieved using perchloric acid in acetone followed by neutralization with ammonium hydroxide to yield **4**. Cyclization of the exocyclic amine was achieved with an aqueous solution of chloroacetaldehyde in mildly acidic sodium acetate buffer at 40° C, yielding the fluorescent product and common intermediate **5**. Substitution of the 2-chloro group in compound **5** using the respective amine yielded the nucleoside products **6a** and **6b**. Synthesis of the carbamate inhibitors **8** and **9** proceeded beginning with nucleoside intermediates **6a** and **6b**. The intermediate nucleoside **6c**, lacking any substitution at the two position was synthesized as previously reported and served as the starting point for compound **10**.^26^ The carbamate backbone was achieved via a two-step one-pot coupling reaction. Addition of *p*-nitrophenyl chloroformate to **6a-c** in pyridine yielded the activated carbonate ester intermediate. Dropwise addition of tryptamine in pyridine to this intermediate produces the penultimate products **7a-c**. Treatment of **7a-c** with triflouroacetic acid in water results in efficient acetonide deprotection, yielding the final products **8-10**. Addition of bulk to the 2-position was detrimental to enzymatic inhibition of hHINT1, as observed via determination of the inhibitor K_i_ by our previously reported continuous fluorescence assay (**Table 1**).^24^ Specifically, the non-substituted ethenoadenosine carbamate **10**, had the lowest K_i_ of the three inhibitors. Addition of the 2-amino group in **8** increased the K_i_ 10-fold, while the 2-methylamino containing compound **9** resulted in a roughly 60-fold increase compared to compound **10**.

**Table 1.**

| Compound | R | K<sub>i</sub> (μM) |
| --- | --- | --- |
| 8 |  | 1.14 ± 0.07 |
| 9 |  | 6.31 ± 0.09 |
| 10 |  | 0.132 ± 0.064 |

Next, we evaluated the effect of ethenoadenosine modifications to the acyl-sulfamate-containing nucleoside inhibitors. Synthesis of the acyl-sulfamate inhibitors was achieved using our lab’s previously reported synthetic route beginning with compounds **6a** and **6b**.^26^ Treatment of compounds **6a,b** with sulfamoyl chloride and triethylamine in DMA yielded compounds **11a,b**. Coupling of **11a,b** with the *N*-hydroxysuccinic acid ester of 3-indole propionic acid (**15**) in the presence of DBU yielded the penultimate products **12a,b.** Deprotection of the acetonide group with aqueous TFA yielded the final products **13** and **14** in good yield. Substitution at the 2-postion did not have the same effect on HINT1 binding for the acyl-sulfamate series as it did for the carbamates. Compound **14**, possessing the N^2^-methyl group, had an inhibition constant roughly half that of **13 (Table 2)**. Replacement of the carbamate backbone of **9**, with the acyl-sulfamate backbone of **14** resulted in a 10-fold improvement in K_i_, a result which we have observed with our previous inhibitors.^4^ However, this improvement was not observed for compounds **8** and **13**, which possess the exocyclic amine at the 2-postion, but lack the N^2^-methyl modification, as these compounds had similar inhibition constants for HINT1.

**Scheme 1.**
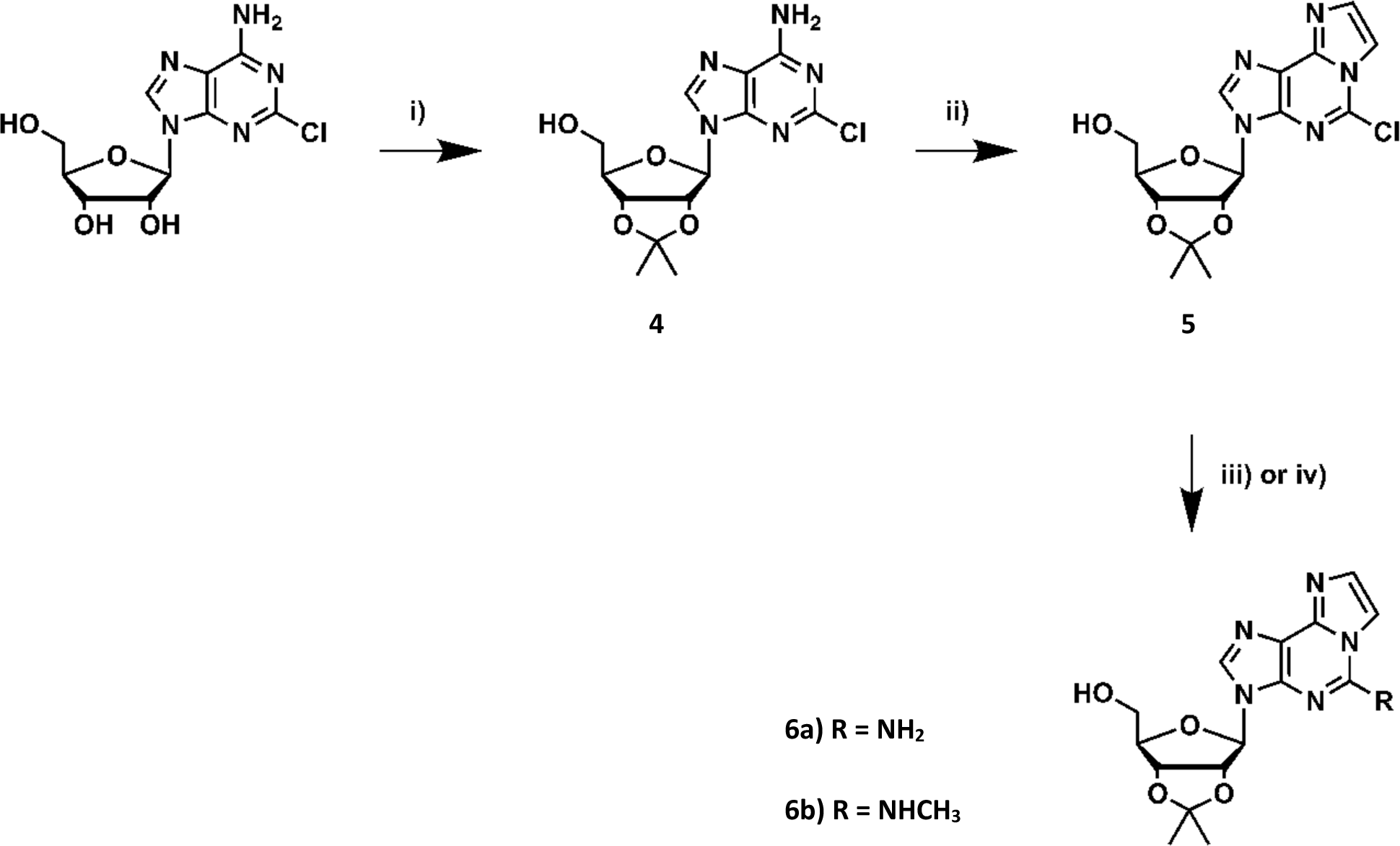
Synthetic Strategy for Preparation of Nucleoside Precursors 6a and 6b. Reagents and conditions: (i) Acetone, perchloric acid, rt, 3 h; (ii) chloroacetaldehyde solution in H_2_O 50% wv, 0.1 M sodium acetate buffer pH 6.5, 40° C, 24 h; iii) 2.0 M NH_3_/Isopropanol, 75° C, sealed tube, on; iv) 2.0 M methylamine/THF, rt, on.

**Scheme 2.**
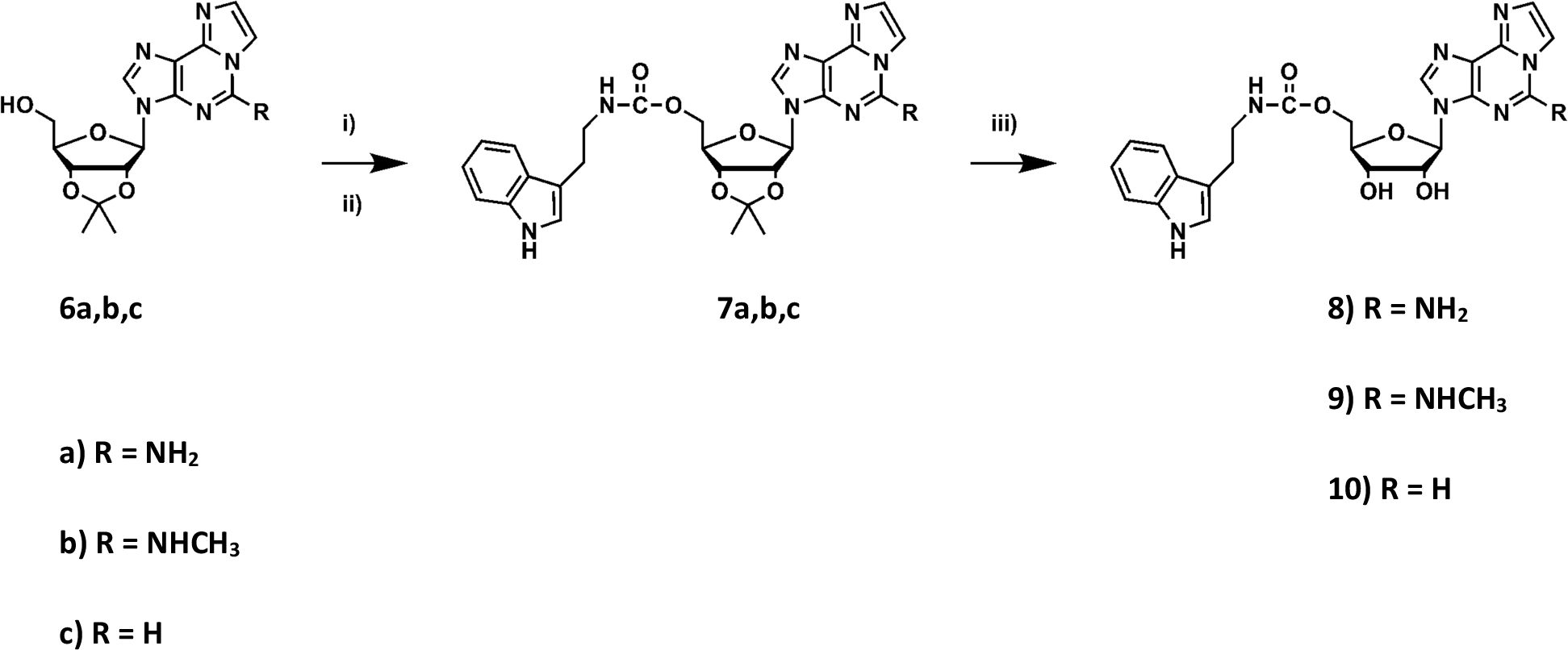
Synthetic Scheme for the Preparation of Target Compounds 8-10. Reagents and conditions: i) p-NO-phenyl chloroformate, pyridine, rt, 3 h; ii) Tryptamine, pyridine, rt, on; iii) 4:1 TFA:H_2_O, rt, 30 min.

**Scheme 3.**
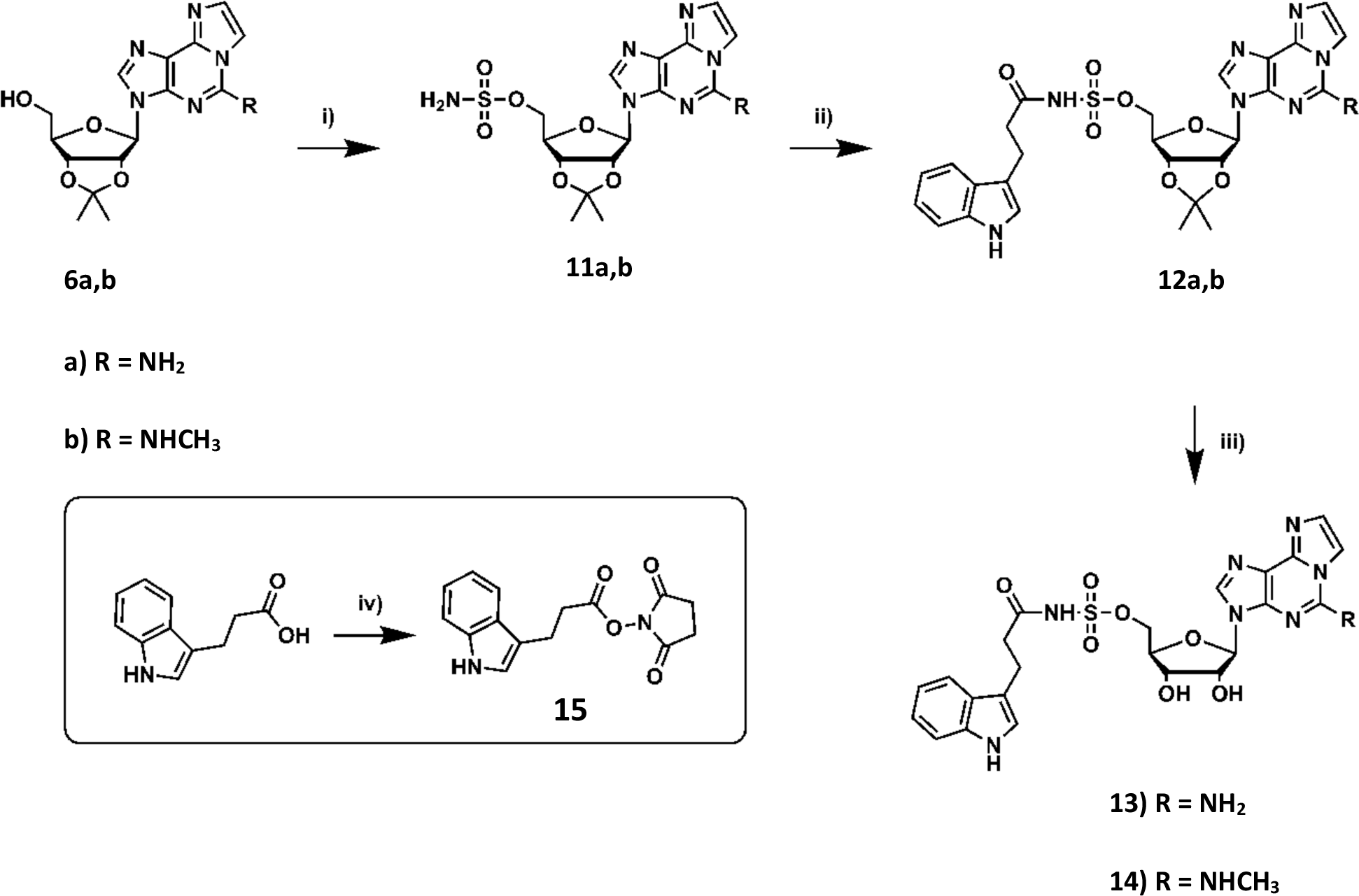
Synthetic Scheme for the Preparation of Target Compounds 13 and 14. Reagents and conditions: i) Sulfamoyl chloride, TEA, DMF, 0° C, 1 h; ii) **15**, DBU, DMF, 0° C, 1 h, rt, on; iii) 4:1 TFA:H_2_O, rt, 30 min; iv) NHS, EDC, THF, rt, 21 h.

**Table 2.**

| Compound | R | K<sub>i</sub> (μM) |
| --- | --- | --- |
| 13 |  | 1.43 ± 0.08 |
| 14 |  | 0.730 ± 0.090 |

### X-Ray Crystallography Reveals the Impact of Ethenoadenosine Base Modifications on Inhibitor Binding to HINT1

To examine the molecular interactions responsible for inhibitor/HINT1 binding, we obtained high resolution X-ray crystallographic structures for compounds **(8-10, 13, 14)** bound to HINT1 (**Figure 2**). Crucially, each of the new inhibitors displayed the typical HINT1 binding position and interactions observed in our previously obtained crystal structures. Notably, the 2’, 3’-OH of each inhibitor ribose forms a tight hydrogen bond with the side chain of Asp43 (2.5-2.7 Å) of HINT1. Further, each inhibitor occupies the hydrophobic binding pocket composed of Ile18, Phe19, Ile22, Phe41, and Ile44. However, substitution of the ethenoadenosine base had a significant impact on the position of each nucleobase for inhibitors **8-10** (**Figure 3A**), with increasing bulk shifting the base further out in the hydrophobic pocket. Compound **10**, lacking any substitution, sat deepest in the binding pocket, while **9**, containing the 2-methylamino substitution was pushed furthest out due the added steric bulk of this modification. Additionally, to accommodate the increased size of **9** its ribose is slightly out of alignment with **8** and **10 (Figure 3A)**, though it still maintains its tight hydrogen bonds with Asp43. In agreement with our hypothesis, the 2-amino group of **8** formed a tight hydrogen bond with the backbone carbonyl of His42 (**Figure 3A,B**). Comparison of the crystal structures of **8** and **TrpGc** reveals that the 2-amino group of each inhibitor participates in a nearly identical hydrogen bonding interaction (**Figure 3B**). Despite this additional hydrogen bond, **10** still binds favorably to hHINT1 compared **8** when looking at their K_i_s, indicating that the additional H-bond does not outweigh the shift in nucleobase position required to accommodate this modification.

**Figure 2.**
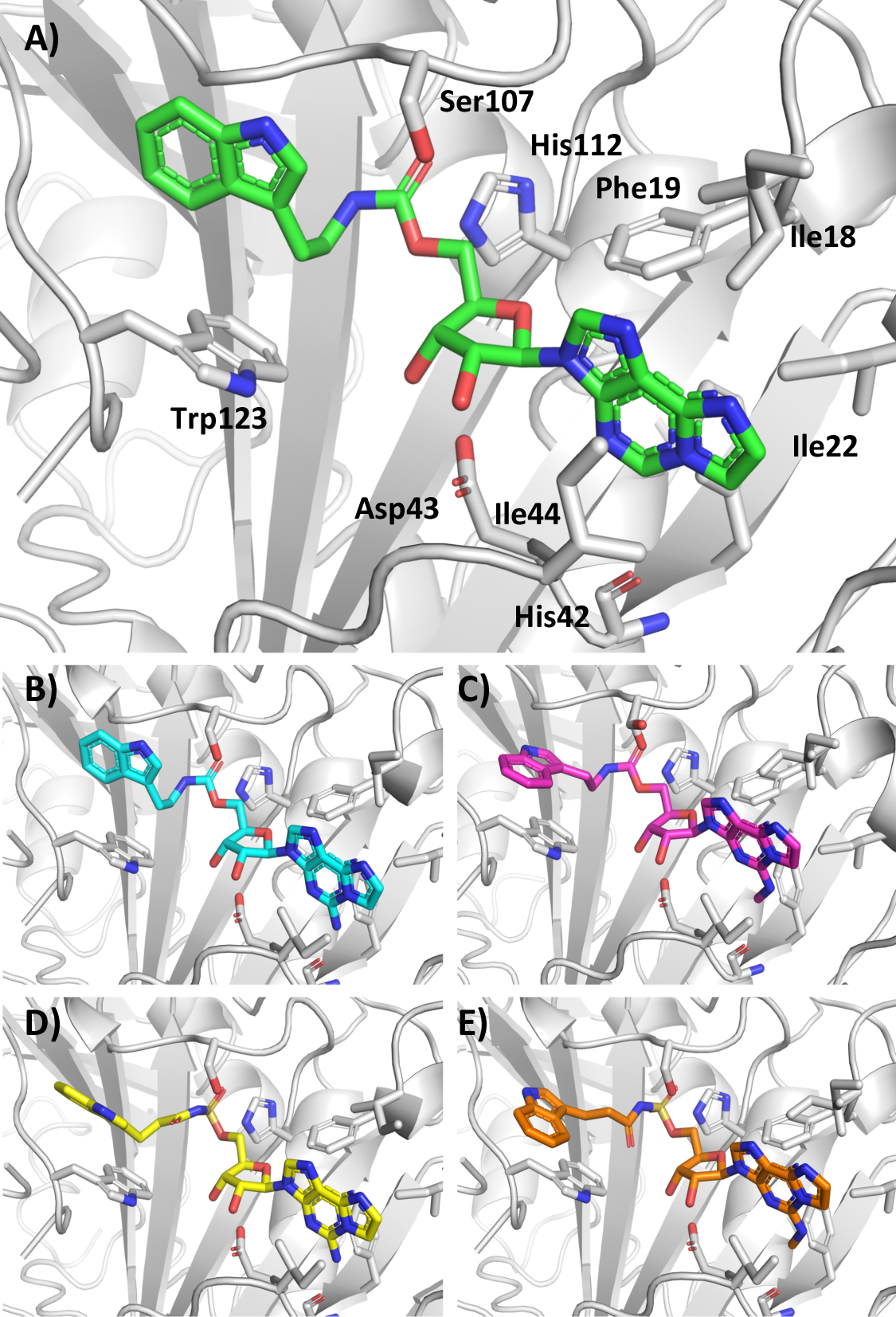
Co-crystal structures of Compounds 8-10, 13, 14 bound to HINT1. A) Compound **10** (Green) bound to HINT1 (gray) depicted as a cartoon. Key residues are labeled. A) Compound **10** (Green). B) Compound **8** (Cyan). C) Compound **9** (Magenta). D) Compound **13** (Yellow). E) Compound **14** (Orange).

**Figure 3.**
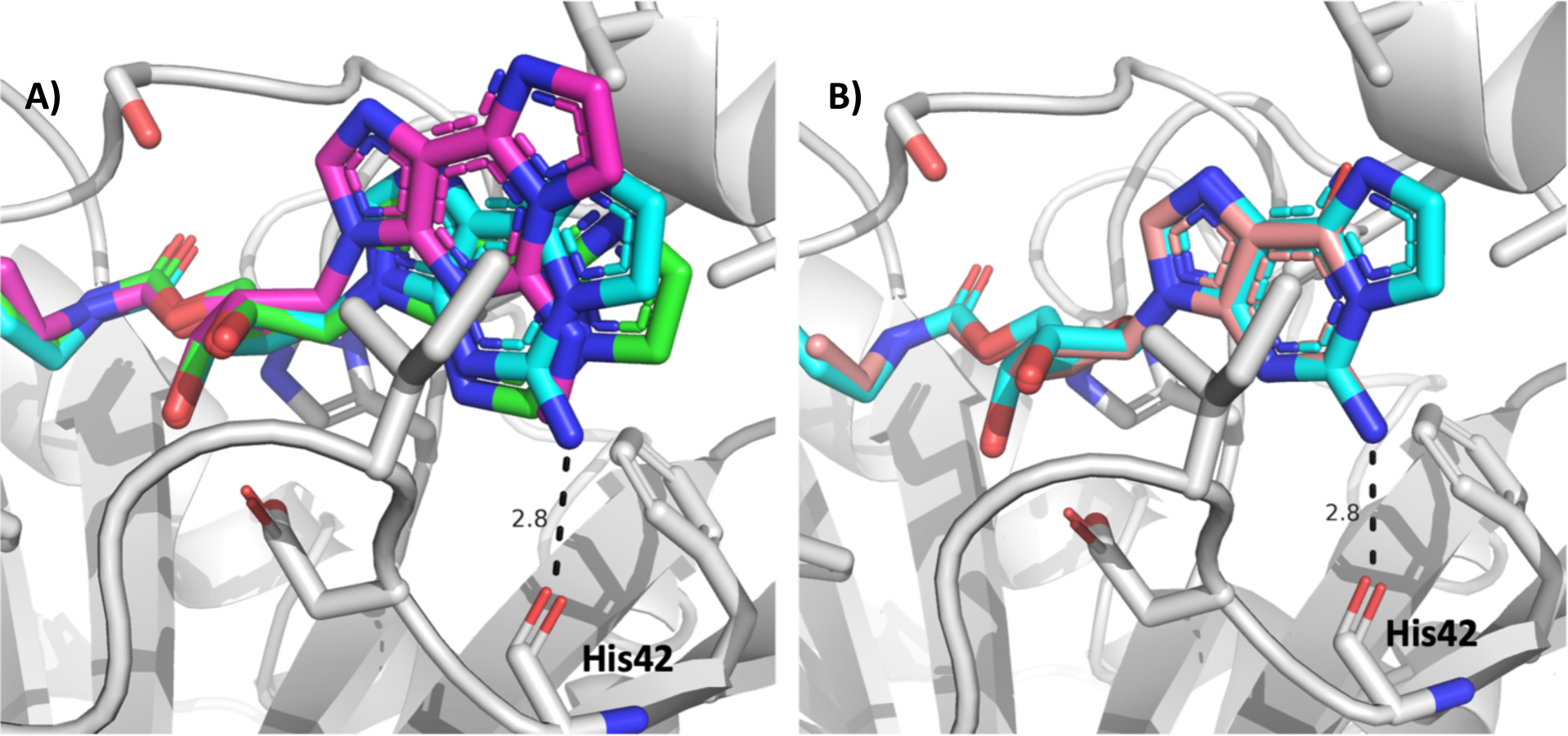
Comparison of Nucleoside Positioning for HINT1 Inhibitors. A) Carbamate Series of HINT1 inhibitors bound to HINT1; Compound **8** (Cyan), **9** (Magenta), and **10** (Green). Increasing nucleobase size results in displacement of the ethenoadenosine base. Compound **8** forms a tight hydrogen bound with the backbone carbonyl of His42. B) Compound **8** (Cyan) and **TrpGc** (Light Pink) (PDB: **6N3V**) bound to HINT1. The 2-aminoethenoadenosine and guanosine nucleobases form identical hydrogen bonds to the backbone carbonyl of His42.

As expected, the position of the carbamate group does not appear to be severely impacted by the alterations made at the ethenoadenosine base, as indicated by both the similar angle of the carbamate groups and the distance between the carbon centers and the nucleophilic nitrogen of His112 (3.1-3.3 Å). However, while the tryptamine side chains of **8** and **10** occupy similar positions, the indole of **9** appears to be rotated roughly 90 degrees, resulting in the indole pointing away from the binding pocket. This is potentially due to the larger size of the nucleobase in **9**. Comparison of the two acyl-sulfamate compounds **13** and **14** reveals a significant change in the backbone geometry between the two inhibitors (**Figure 4**). The acyl sulfamate of **13** displays a similar binding mode to our previously reported **3**, in which the side chain acyl group participates in a hydrogen bond with Ser107 (2.9 Å). By comparison, the side chain acyl group of **14** is rotated roughly 90 degrees away from the side chain of Ser107. As a result, the distance between the acyl group and Ser107 is too far for a potential hydrogen bond (3.8 Å). However, the absence of this hydrogen bond does not appear detrimental to HINT1 binding for **14**, as its K_i_ is roughly two-fold lower than **13** (**Table 2**). The sulfamate inhibitors also display differences in tryptamine position, with the indole groups of **13** and **14** flipped 180° from each other (**Figure 4**). The differences inside chain position, especially the solvent-accessible indole, could contribute to their unique pharmacology *in vivo*.

**Figure 4.**
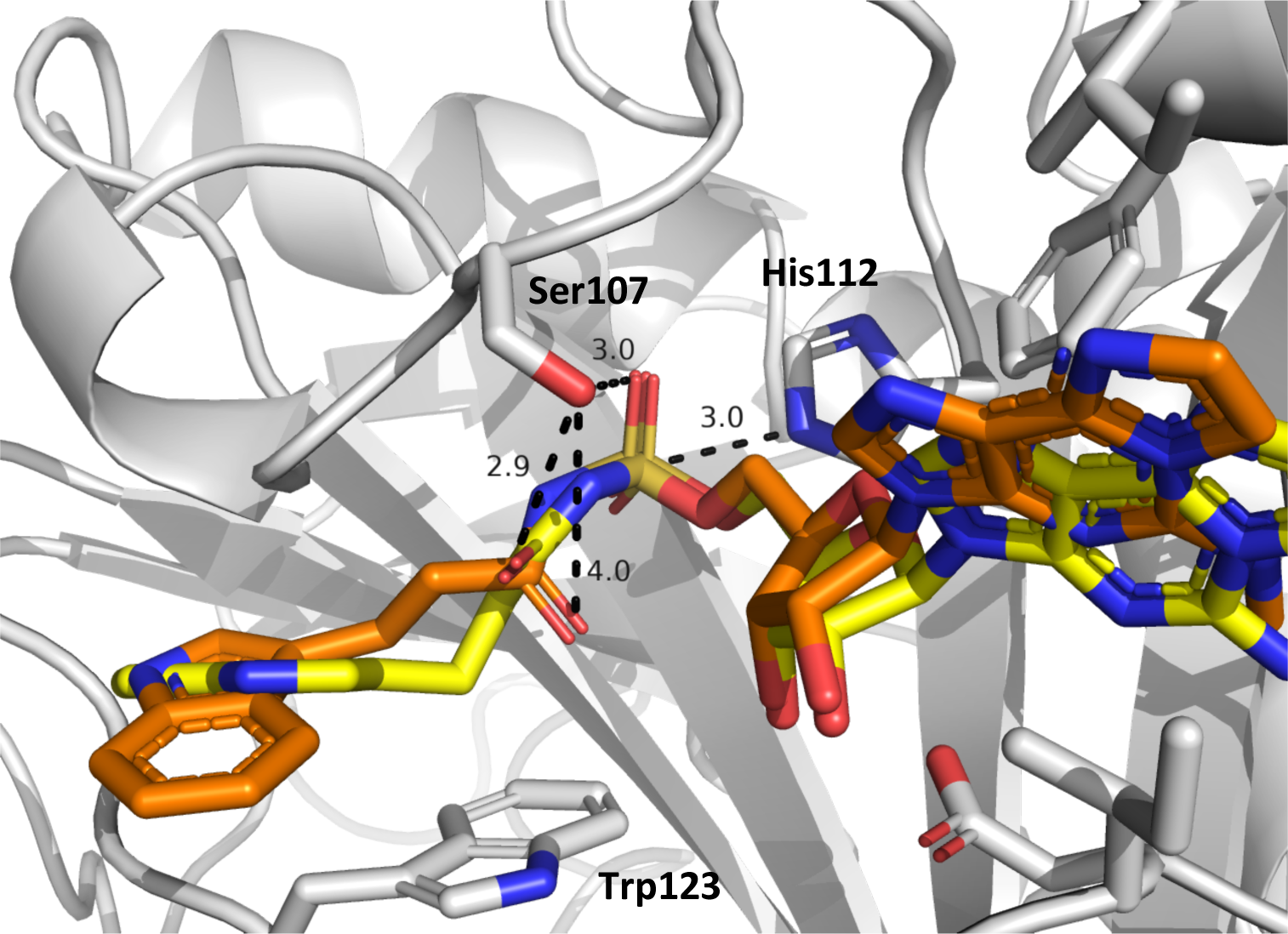
Comparison of Acyl-Sulfamate Side Chains for Compounds 13 and 14 Bound to HINT1. The acyl-sulfamate side chains of compound **13** (Yellow) and **14** (orange) display different geometries. The acyl group of **13** makes a tight hydrogen bond to Ser107 (2.9 Å), while the side chain of **14** is rotated 90°, no longer in position to form a hydrogen bond. The indoles of **13** and **14** are rotated 180° from each other. The sulfamate groups of both inhibitors are in similar positions, with the sulfamate group in close position to active site His112 and hydrogen bonding to Ser107.

### TrpGc Blocks Morphine’s Inhibition of NMDA-Evoked Behaviors and Thermal Hyperalgesia

As previously described, NMDA delivered to the intrathecal space of the spinal cord produces a set of quantifiable nociceptive behaviors in mice: first, a transient, caudally-directed set of scratching and biting behaviors and second, a transient thermal hyperalgesia that is observed using the warm water tail flick assay.^27–28^ Both the behavioral scratching and the thermal hyperalgesia can be inhibited by delivery of morphine prior to NMDA (**Figure 5**). This response is calculated as the percent maximum possible effect (%MPE). We first assessed whether the HINT1 inhibitor compounds were effective at reducing morphine’s efficacy at reducing the NMDA-induced scratching and biting behaviors. While **TrpGc** effectively reduced morphine’s efficacy in this assay, consistent with previous reports, none of the newly developed compounds showed similar efficacy (**Figure 5A**).^4^ The observed ED_50_ for TrpGc was 0.56nmol (0.21 – 1.4nmol, CI), consistent with our previous report.^4^ The ED50 could not be calculated for any other tested compound due to the lack of efficacy.

**Figure 5.**
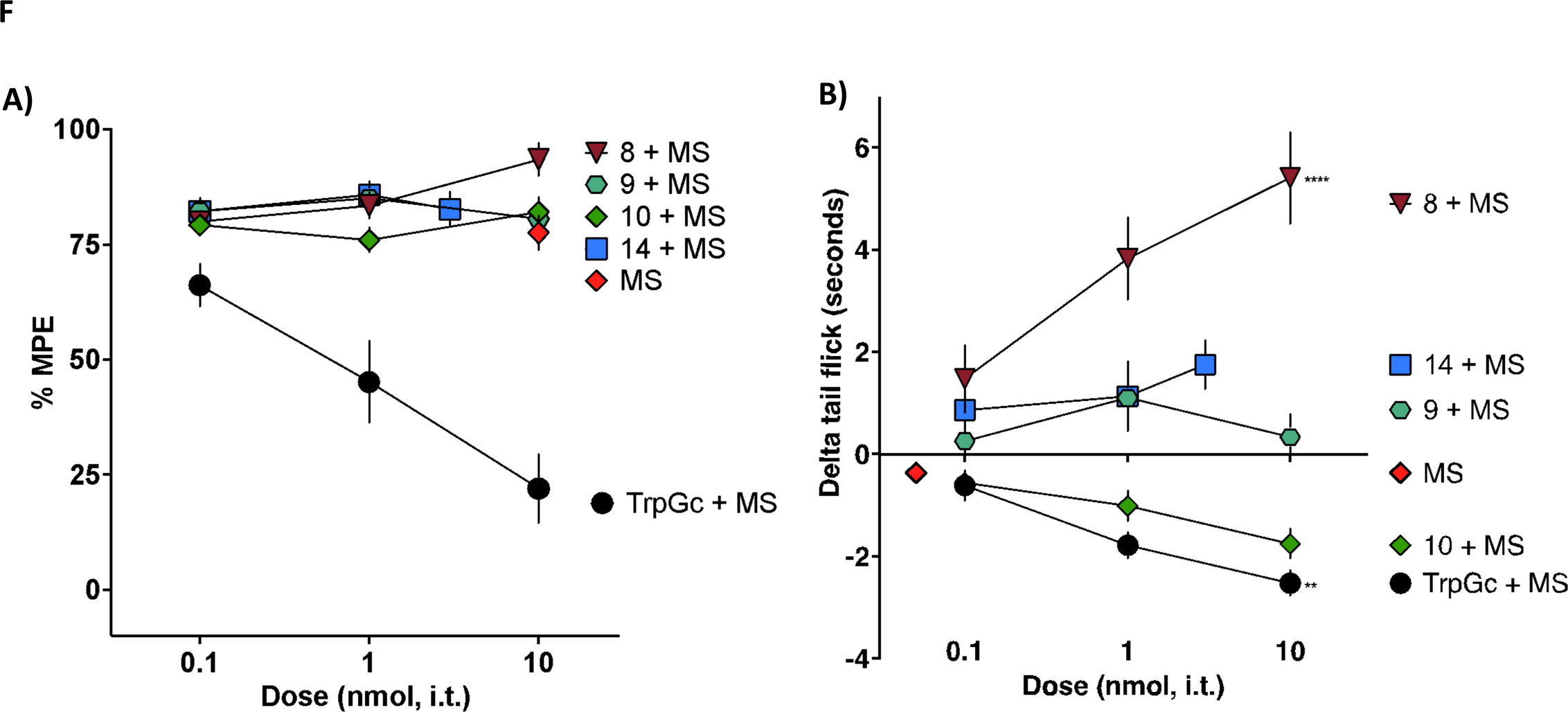
HINT1 Inhibitors Effect on Morphine’s Inhibition of NMDA-Evoked Nociceptive Behaviors. Male and female mice were given intrathecal treatment of either morphine sulfate (MS) or MS + a HINT1 inhibitor. They were then given an intrathecal injection of NMDA and scratching and biting behaviors were counted and transient thermal hypersensitivity was measured by tail flick assay in a 49°C water bath.. N = 6/drug/dose, M + F. (A) TrpGc alone inhibited MS’s inhibition of scratching and biting behaviors, with an observed ED50 of 0.56 nmol (0.2-1.4 nmol, CI). ED50s could not be calculated for any other compound due to a lack of a dose-related effect in this assay. (B) Pretreatment with TrpGc (black circles) significantly attenuated MS-mediated inhibition of NMDA-evoked transient thermal hypersensitivity, whereas pre-treatment with compound 8 enhanced MS-mediated prevention of the development of thermal hypersensitivity following intrathecally-delivered NMDA. **p < 0.01, **** p < 0.0001. One-way ANOVA with Dunnett’s posthoc test with multiple comparisons to a control (MS delivered alone).

Following assessment of the scratching and biting behaviors, the same subjects were then tested for the presence of transient thermal hyperalgesia in the warm water tail flick assay, and their responses were compared to a pre-NMDA baseline tail flick. While subjects who receive only NMDA show a decrease in their tail flick, termed delta tail flick, subjects pre-treated with morphine do not display this transient thermal hypersensitivity (**Figure 5B)**. **TrpGc** again significantly inhibited morphine’s efficacy in preventing this thermal hypersensitivity, with **10** showing a similar effect, but the 10 nmol dose was not statistically significantly different from morphine alone. Compounds **9** and **14** had little impact on morphine’s efficacy in this assay, whereas compound **8** co-administered with morphine showed a marked and significant decrease in thermal hypersensitivity as measured by an increase in tail flick latency as compared to morphine alone. This result suggested either enhanced morphine efficacy or had a direct analgesic effect in this assay.

### TrpGc and 9 Inhibit Endomorphin-2 Tolerance

Endomorphin-2, an endogenous MOR agonist, produces acute tolerance when delivered to the intrathecal space at a high dose.^29^ Male and female mice were pre-treated with either vehicle control or a tolerance-inducing dose of endomorphin-2 (10 nmol). Thirty minutes following this first injection, tail flick latencies return to baseline levels and a second injection of endomorphin-2 (10 nmol) was administered. Following this second injection, tail flick latencies are again assessed. Mice that received a vehicle pre-treatment demonstrated full antinociception following the second injection, the probe dose of endomorphin-2. However, mice pre-treated with endomorphin-2 showed reduced antinociception following the probe dose, a demonstration of acute spinal tolerance. To determine whether the inhibitor compounds could effectively inhibit the development of acute spinal tolerance, mice were pre-treated with HINT1 inhibitor. **TrpGc** and **9** reduced the magnitude of acute spinal analgesic tolerance, whereas **13** and **14** displayed potentiated tolerance in this assay (**Figure 6**).

**Figure 6.**
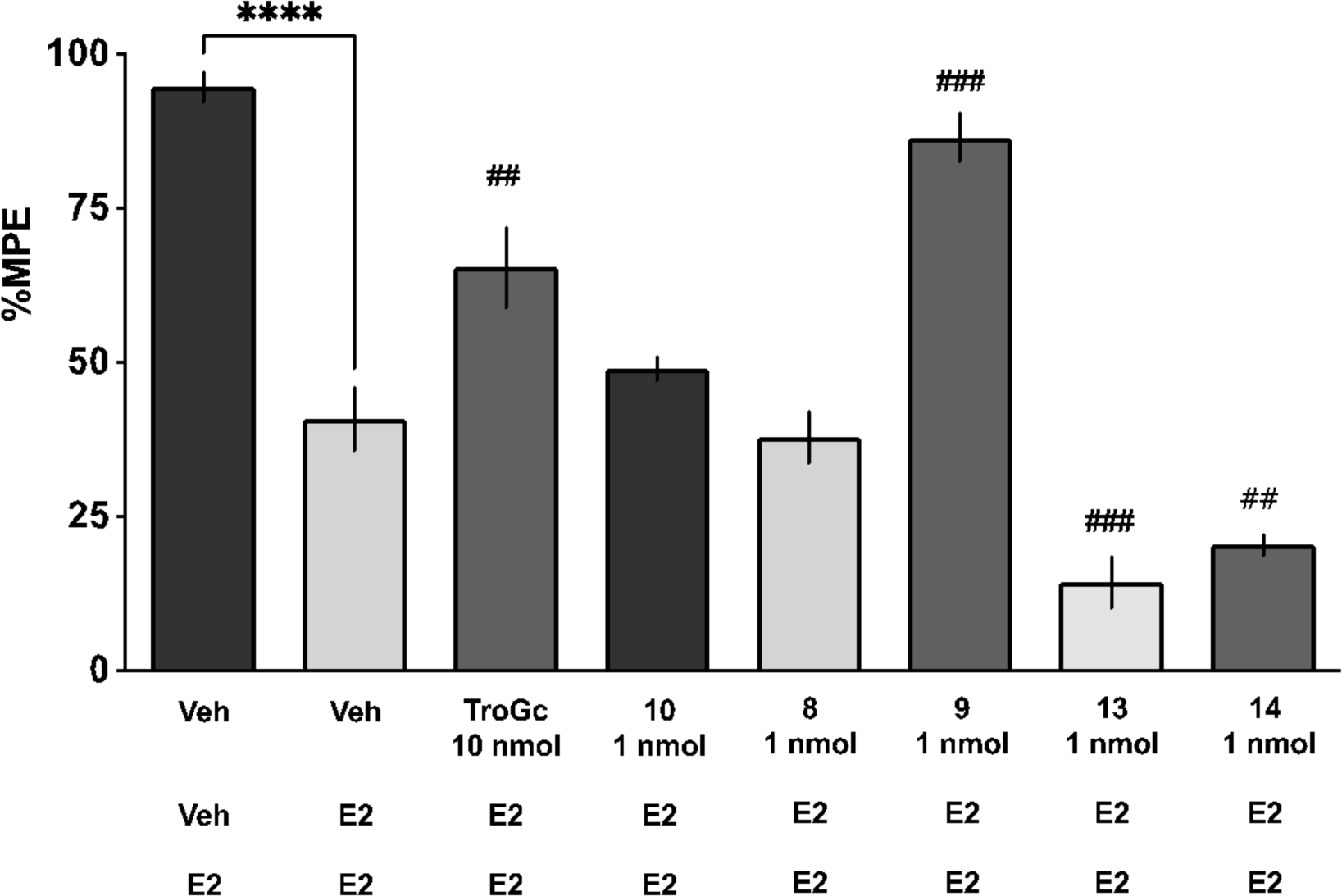
TrpGc and 9 inhibit endomorphin-2 tolerance. The spinal antinociceptive effect of endomorphin-2 is significantly greater in male and female mice pretreated with vehicle (leftmost column) as compared with mice pretreated with endomorphin-2 (2^nd^ column), indicating an ultrarapid development of tolerance as assayed by a warm water bath tail flick (52.5°C), as previously described. Treatment with TrpGc prior to the first endo-2 injection can prevent this development of tolerance. The TrpGc analogues 506.52 and 641.72 were also able to prevent the development of endo-2 tolerance. ****indicates significant difference from saline-endo-2 control by Student’s t-test, p < 0.001. # indicates significant difference from pre-treatment with endomorphin-2 alone (2^nd^ column) by one-way ANOVA with Dunnett’s posthoc test with multiple comparisons to a control. N = 6/group, M + F.

### Intrathecal Administration of 8, but No Other Inhibitor Leads to the Development of Analgesia

To assess the analgesic efficacy of the line of Hint1 inhibitor compounds as single agents rather than in combination with morphine, mice were intrathecally injected with increasing doses of the HINT1 inhibitor and their responses in the warm water tail flick assay were evaluated. Once again, **8** showed an analgesic effect whereas the rest of the Hint1 compounds did not show analgesic efficacy in this assay (**Figure 7**). To determine if the observed response was due to a sedative or anesthetic effect, we carried out a series of motor studies. The 10 nmol of the inhibitors were dosed intrathecally and the animals tested 5 min afterwards. As can be seen in **Figs. S6 and S7**, no difference was observed between saline and any inhibitor for the rotarod motor assay as well as the open field assay. Thus, these data support the observation of analgesia seen with the tail flick assay for compound **8**.

**Figure 7.**
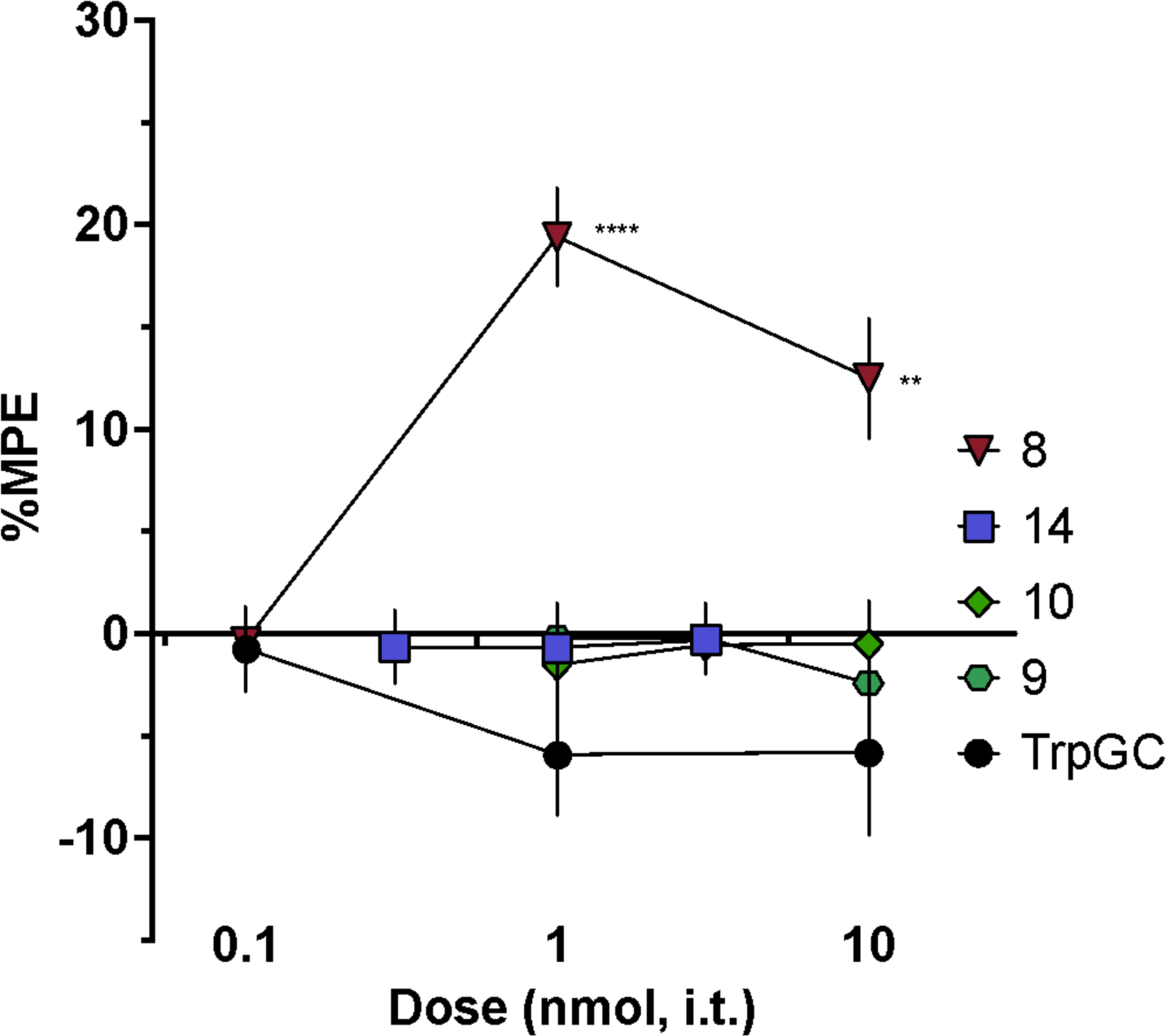
Analysis of Analgesia Following Intrathecal Dosing of HINT1 Inhibitors. Assessment of Hint1 inhibitors in warm water tail flick assay (52.5°C). Following baseline tail flick assessment, subjects were given intrathecal injections of increasing doses of TrpGc or Hint1 analogue compound, and tail flick latency was again assessed. Responses are reported as %MPE, which was used to generate dose-response curves. Compound 8 showed increasing MPE at 1 and 10 nmol, i.t. doses as compared to 0.1 nmol, while the other compounds did not have a dose-related effect. **p < 0.01, **** p < 0.0001. One-way ANOVA with Dunnett’s posthoc test with multiple comparisons to a control (lowest dose administered).

### Treatment of cells with HINT1 Inhibitors does not lead to loss of HINT1 or p53

Previously, knock out studies of HINT1 were shown to lead to the potential for enhanced tumorgenicity.^30^ This effect was later associated with the loss of p53 expression.^25, 31^ The loss of p53 expression could be rescued by the introduction of wild type HINT1, as well as the H112N catalytically dead active site mutant.^25^ Recently, we have shown that treatment of cells with HINT1 inhibitors can block the activation of antiviral proTides by HINT1.^32^ Consequently, we chose to determine if treatment of cells with HINT1 a inhibitor affected the expression levels of HINT1 and p53 in cells. The breast cancer cell line BT549^(HINT1+, p53+)^ and neuroblastoma cell line, SH-SY5Y^(HINT1+, p53-)^, were treated for 48 h with up to 100 uM of variable concentrations of inhibitor, Trp-Gc. No cytotoxicity was observed, with the cells growing at the same rate as non-treatment cells. Moreover, by western blot analysis, compared to the control non-treatment cells, no significant difference in the amount of HINT1 for either cell line was observed and no change in the amount of p53 expression was observed for BT549 cells (**Fig. 8, Fig. S9**). Thus, consistent with the finding that inactivating active site mutations of HINT1 do not effect p53 expression, inhibitors of HINT1 do not affect either HINT1 or p53 expression, nor have an effect on cellular viability or growth.^25^

**Figure 8.**
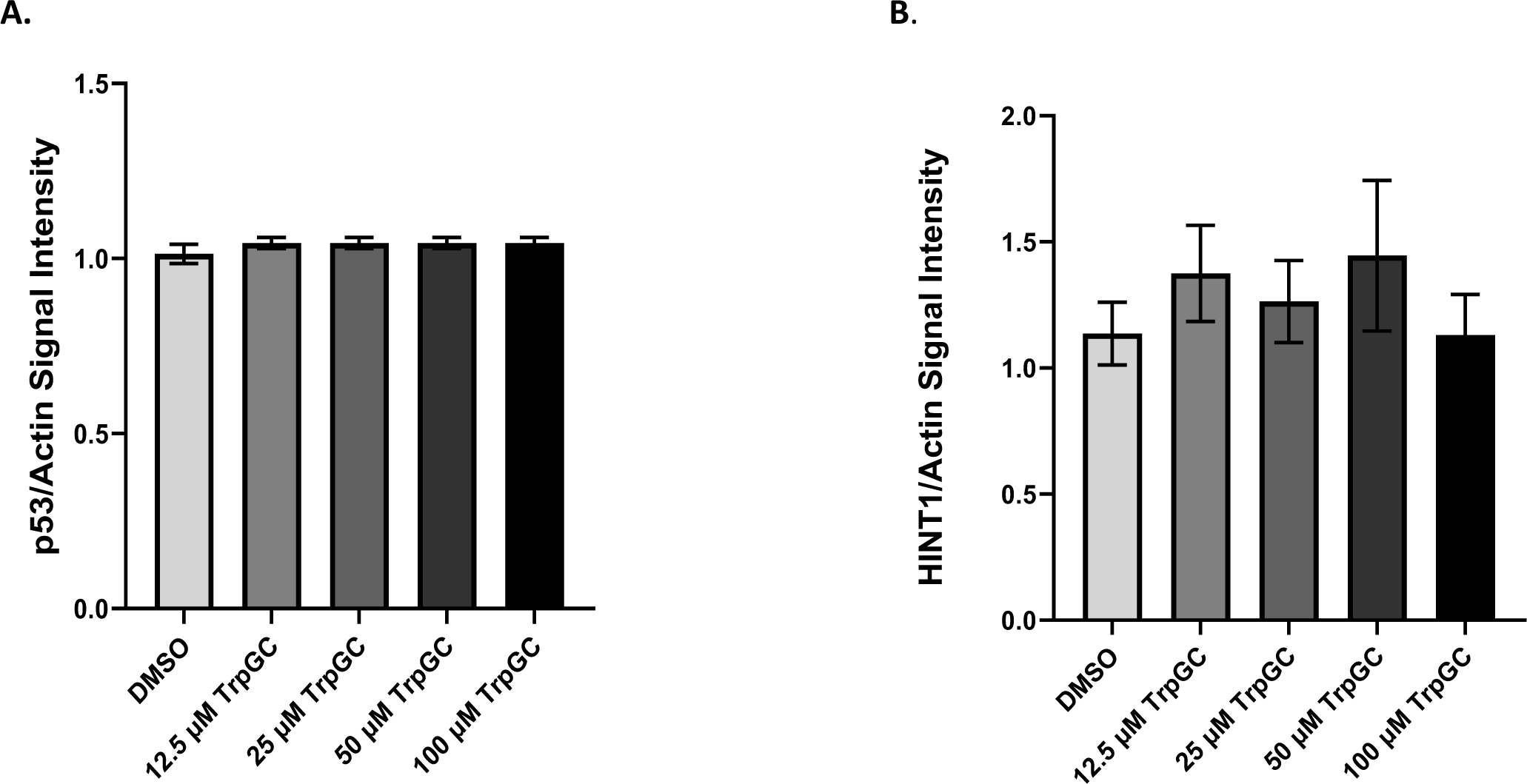
Treatment of BT549^(HINT1+, p53+)^ cells with HINT1 Inhibitor has no effect on HINT1 or p53 expression. BT549 cells were treated with the HINT1 inhibitor, TrpGc, for 72 h, followed by western blot analysis. Western blot densitometry analysis was performed using ImageJ, and relative a) p53 and b) HINT1 band density ratio to β-actin control were determined. Statistical analysis was performed using Brown-Forsythe and Welch ANOVA followed by Dunnett’s T3 multiple comparisons test. Data shown as mean ± SEM (n=5 individual experiments). No significant difference among treatment means was determined for both proteins (p>0.05). Representative western blot images can be found at **Fig. S8** and **Fig. S9**.

### Conclusions

Previous work by our laboratory demonstrated the exciting potential for small molecule HINT1 inhibitors to affect various pain pathways.^4, 23^ Based on the tight binding and unique pharmacological effect of **3**, our lab sought to develop ethenoadenosine based analogues with the goal of improving HINT1 enzymatic inhibition and furthering our understanding of the role of HINT1 in the interplay of MOR and NMDAR. *In vitro*, we confirmed our hypothesis that addition of the 2-amino group to the ethenoadenosine nucleobase (compounds **8** and **13**) results in the formation of a hydrogen bond with the backbone carbonyl of His42, identical to that of the guanosine-based inhibitor **TrpGc**. Despite this, compound **10**, which lacks the amine at the 2-postion, was the strongest inhibitor of HINT1 enzymatic activity as observed by our continuous fluorescence assay. Further, addition of the N^2^-methyl group to the exocyclic amine (compounds **9** and **14**) disrupts this hydrogen bond and shifts the nucleobase further up in the hydrophobic pocket. Interestingly, while this modification resulted in decreased binding for the carbamate inhibitors (Compound **8** and **9**), the addition of the N^6^-methyl group improved binding for the acyl-sulfamates (Compounds **13** and **14**).

*In vivo*, we evaluated the effect of HINT1 inhibitors on MOR-NMDAR crosstalk via their impact on MOR inhibition of NMDAR activation and the development of acute endomorphin-2 tolerance. We observed that minor modifications to the ethenoadenosine scaffold resulted in major changes in their activity. Consistent with our previous results, **TrpGc** displayed broad activity, preventing morphine from blocking NMDA evoked transient thermal hypersensitivity and behaviors as well as preventing the development of endomorphin-2 tolerance.^4^ Replacement of the guanosine base with the tricyclic ethenoadenosine base **(10)** abolished activity observed in all three assays. Addition of a N^2^-methyl group **(9)** to the ethenoadenosine base resulted in ablation of its activity against MOR inhibition of NMDA evoked transient thermal sensitivity and behaviors but was observed to prevent the development of acute endomophin-2 tolerance. Perhaps the most intriguing result came from **(8)**, which displayed enhancement of morphine efficacy in the NMDA evoked transient thermal hypersensitivity assay, as opposed to inhibition that we have observed with the other HINT1 inhibitors. Further, **(8)** was observed to induce analgesia following intrathecal injection, something not yet observed with any HINT1 inhibitor. Lastly, replacement of the carbamate backbone of compounds **(8)** and **(9)** with an acyl-sulfamate, compounds **(13)** and **(14),** seems to largely ablate their activity *in vivo*, with no efficacy observed in the NMDA evoked transient thermal hypersensitivity and behavior assays. This could potentially be explained by the large difference in polarity between the two sets of compounds. Interestingly, treatment with these compounds resulted in increased development of endomorphin-2 tolerance, though the mechanism behind this activity is not clear.

The SAR efforts of this work have resulted in a set of HINT1 inhibitors that selectively impact unique pathways of MOR-NMDAR crosstalk *in vivo*. Analysis of the *in vitro* and *in vivo* results has yet to establish a clear connection between these two activities. Consistent with our previous works, HINT1 binding affinity was not correlated with *in vivo* activity. Evaluation of the co-crystal structures for the HINT1 bound inhibitors reveals that despite the presence of similar hallmark interactions, alterations to the nucleobase have a profound effect on the positioning of both the nucleobase and the inhibitor side chain. Due to the solvent accessible nature of these regions, each HINT1 inhibitor alters the molecular surface of the HINT1-Inhibitor complex (**Figure 9)**. Because HINT1’s involvement in MOR-NMDAR crosstalk relies on its participation in several protein-protein interactions, alteration to its molecular surface could have a significant impact on its ability to participate in these key interactions. Interestingly, due to the unique pharmacological effects observed for these inhibitors *in vivo*, it appears that small-molecule intervention can potentially alter HINT1’s interactions with some specificity, resulting in the selective inhibition of one pathway versus another, though further studies into the specific effects on these protein-protein interactions are needed. Additionally, we observed that **8** produces analgesia following intrathecal administration, though the mechanism behind this unexpected activity remains to be determined. Furthermore, we have demonstrated that effects on p53 expression that maybe tied to HINT1 expression are not observed for cells treated with HINT1 inhibitors. Together, these results highlight the intriguing role of HINT1 in MOR-NMDAR crosstalk and the pharmacological possibilities that small molecule inhibition of HINT1 can afford. Evaluation of the mechanisms behind these pathways at the molecular level represents the crucial next step in advancing our understanding of the role of HINT1 in the CNS and its potential as a novel target for pain therapeutic development.

**Figure 9.**
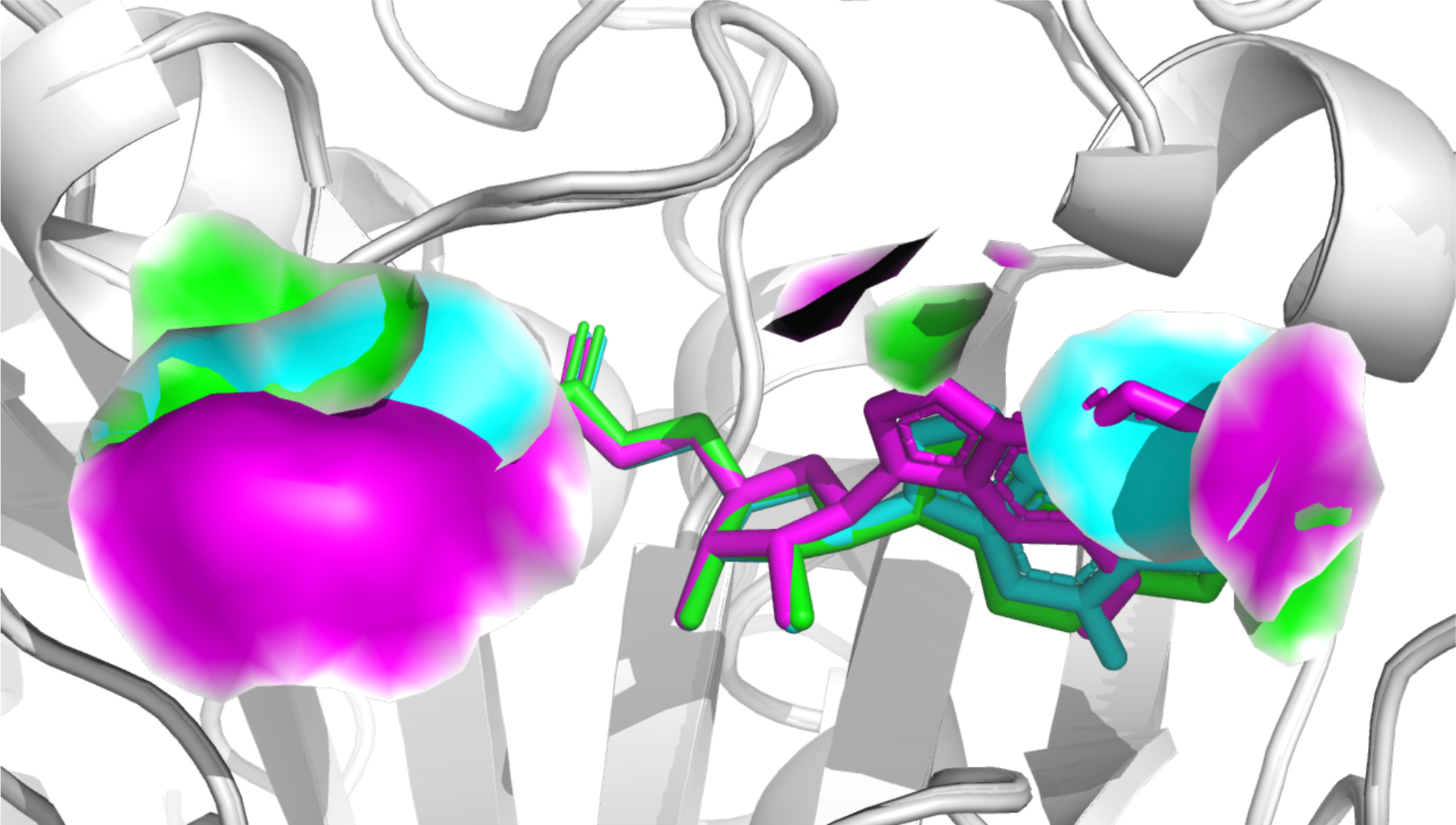
Comparison of the molecular surface of carbamate-based nucleoside inhibitors. Compounds 8 (**Cyan**), 9 (**Magenta**), and 10 (**Green**) bound to HINT1. The contributions of the inhibitors to the molecular surface of the bound HINT1/inhibitor complex are displayed in their respective colors. Changes to the molecular surface occur at the nucleobase and tryptamine side-chain.

## Materials and Methods

### General Methods

All chemicals and reagents were obtained from commercial sources and were used without further purification. Anhydrous *N*,*N*-Dimethylformamide (DMF) was obtained from a dry solvent purification system (MBraun) and dispensed under argon. All reactions were performed under an atmosphere of dry nitrogen unless otherwise noted. All silica gel chromatography and preparative reverse phase purification was performed on a Teledyne Isco CombiFlash Rf system, using Redisep Rf high performance gold silica gel columns (for normal phase purifications) and Redisep Rf high performance gold C18 columns (for reverse phase purifications). Reverse phase purifications were with water and acetonitrile. Lyophilization of compounds after reverse phase purification was performed on a FreeZone 12 Plus freeze dry system (Labconco). All analytical HPLC analyses were performed on a Beckman Coulter System Gold instrument using a Haisil 100 C18 column (4.6 mm X 250 mm, 5 μm, Higgins Analytical Inc.); eluting with a gradient of 10% acetonitrile in phosphate buffer (10 mM or 50 mM, pH 7.4) to 90% acetonitrile in phosphate buffer over 24 minutes at a flow rate of 1.0 mL/min. All Nuclear Magnetic Resonance spectra were obtained on a Brucker Avance III HD 500 MHz spectrometer at ambient temperature. Chemical shifts were recorded in parts per million using the solvent peaks as internal references. All high-resolution mass spectroscopy (HRMS) were performed on a LTQ Orbitrap Velos instrument (Thermo Scientific) in positive-ion mode.

#### Synthesis of 2’,3’-Isopropylidine 2-chloroadenosine (4)

To a round bottom flask was added 2-cl-adenosine with acetone. To the flask was added perchloric acid dropwise. The reaction stirred at room temperature for 3 hours. The reaction was neutralized with ammonium hydroxide ∼5 ml. The reaction was concentrated under reduced pressure and purified using normal phase chromatography (0-15% MeOH/DCM). Relevant fractions were pooled and concentrated under reduced pressure. Yield: quant. The ^1^H NMR spectrum was (DMSO-d_6_): 1.34 (s, 3H), 1.55 (s, 3H), 3.54 (td, 2H), 4.22 (td, 1H), 4.94 (dd, 1H), 5.08 (d, 1H), 5.29 (dd, 1H), 6.06 (d, 1H), 7.86 (s, 2H), 8.36 (s, 1H). ^13^C-DMSO-d_6_: ^13^C NMR (126 MHz, DMSO) δ 25.68, 27.51, 61.95, 81.73, 83.92, 87.16, 89.84, 113.57, 118.58, 140.42, 150.44, 153.56, 157.29. ESI-MS [M+H] 342.2.

#### Synthesis of 2’,3’-Isopropylidine 2-Chloroethenoadenosine (5)

MD-3-17 (2.00 g, 5.85 mmol) was added to a round bottom flask with 50% w/v chloroacetaldehyde/H_2_O (36.00 ml, and 0.1 M NaOAc. The pH was adjusted to 6.5 using 1 M HCl. The reaction was stirred at 40°C for 48 hours. The reaction was concentrated under reduced pressure, then extracted with ethyl acetate and bicarb. The organic layer was collected and purified using normal phase flash chromatography (0-7% MeOH/DCM). The relevant fractions were pooled and concentrated under reduced pressure. Yield: 871 mg, 40.7%. The ^1^H NMR spectrum was (DMSO-d_6_): 1.35 (s, 3H), 1.57 (s, 3H), 3.57 (td, 2H), 4.27 (m, 1H), 5.00 (dd, 1H), 5.09 (t, 1H), 5.36 (dd, 1H), 6.24 (d, 1H), 7.69 (d, 1H), 8.09 (d, 1H), 8.59 (s, 1H). ^13^C-DMSO-d_6_: δ 141.77, 140.79, 138.19, 133.63, 133.52, 123.00, 113.65, 113.34, 90.36, 87.44, 84.33,

#### Synthesis of 2’,3’-Isopropylidine 2-aminoethenoadenosine (6a)

To a sealed tube was added MD-3-29 (300 mg, 0.82 mmol) and 2.0 M NH3/Isopropanol (6.72 ml, 13.44 mmol). The flask was sealed and heated to 75°C overnight. The crude mixture was transferred to a round bottom flask and concentrated under reduced pressure. The product was purified using normal phase chromatography (0-15% MeOH/DCM). Relevant fractions were pooled and concentrated under reduced pressure. Yield: 190 mg, 67%. The ^1^H NMR spectrum was (DMSO-d_6_): δ 1.35 (s, 3H), 1.55 (s, 3H), 3.55 (qt, 2H), 4.16 (td, 1H), 5.05 (m, 2H), 5.32 (dd, 1H), 6.12 (d, 1H), 7.43 (d, 1H), 7.62 (s, 2H), 7.94 (d, 1H), 8.12 (s, 1H). ^13^C-DMSO-d_6_: δ 25.76, 27.57, 62.14, 81.78, 84.19, 87.30, 89.21, 110.08, 113.53, 116.47, 132.14, 136.42, 140.70, 142.34, 145.48. ESI-MS [M+H] 347.2.

#### 2’,3’-Isopropylidine N^2^-methyl-2-aminoethenoadenosine (6b)

To a dry round bottom flask was added MD-3-9 (300 mg, 0.56 mmol) and 2.0 M methylamine/THF (7 ml, 14.00 mmol). The flask was purged under nitrogen gas/vacuum and stirred at room temperature overnight. The product crashed out as a white precipitate, the reaction mixture becoming thick. Volatiles were removed under reduced pressure. The product was purified by normal phase chromatography (0-15% MeOH/DCM). The relevant fractions were pooled and concentrated under reduced pressure. Yield: 223 mg, 75% The ^1^H NMR spectrum was (DMSO-d_6_): 1.35 (s, 3H), 1.56 (s, 3H), 3.06 (d, 3H), 3.55 (m, 2H), 4.16 (m, 1H), 4.98 (t, 1H), 5.05 (dd, 1H), 5.47 (dd, 1H), 6.19 (d, 1H), 7.44 (d, 1H), 7.87 (q, 1H), 7.95 (d, 1H), 8.10 (s, 1H). ^13^C-DMSO-d_6_: 144.96, 142.22, 140.39, 137.13, 132.19, 116.55, 113.49, 109.53, 89.48, 87.15, 83.84, 81.93, 62.02, 28.68, 27.51, and 25.16. ESI-MS [M+H] 361.3.

#### 2’,3’-Isopropylidine-5’-O-[(3-Indolyl)-1-Ethyl]Carbamoyl 2-aminoethenoadenosine (7a)

MD-3-39 (200 mg, 0.58 mmol) and 4-nitrophenyl chloroformate (186 mg, 0.92 mmol) were added to a round bottom flask and dissolved in pyridine (8 ml). The reaction was stirred for 2.5 hr at room temperature. To the reaction mixture was added tryptamine (185 mg, 1.15 mmol) dissolved in pyridine(2 ml). The reaction was stirred overnight at RT. The reaction was concentrated under reduced pressure. The crude mixture was taken up in ethyl acetate and sodium bicarbonate. The organic layer was extracted twice and combined. The organic layer was washed with bicarb, brine and water. The organic layer was collected, dried over MgSO4, filtered, and dried under reduced pressure. The product was purified using flash chromatography (0-15% DCM/MeOH). The relevant fractions were pooled and concentrated under reduced pressure. Yield: 74.5 mg, 24.2%. The ^1^H NMR spectrum was (DMSO-d_6_): 1.35 (s, 3H), 1.56 (s, 3H), 2.81 (t, 2H), 3.18 (d, 1H), 3.23 (dd, 2H), 4.30 (m, 2H), 5.18 (dd, 1H), 5.38 (dd, 1H), 6.18 (d, 1H), 6.97 (m, 1H), 7.05 (m, 1H), 7.14 (d, 1H), 7.32 (d, 2H), 7.37 (t, 1H), 7.44 (d, 1H), 7.50 (d, 1H), 7.66 (s, 2H), 7.95 (d, 1H), and 8.09 (s, 1H). ^13^C-DMSO-d_6:_ 156.24, 145.52, 142.32, 140.54, 136.67, 136.61, 132.16, 127.65, 123.10, 121.35, 118.68, 118.64, 116.62, 113.79, 112.03, 118.81, 110.11, 89.14, 84.81, 84.08, 81.79, 64.61, 49.07, 27.57, 25.89, and 25.80. ESI-MS [M+H] 533.3.

#### Synthesis of 2’,3’-Isopropylidine-5’-O-[(3-Indolyl)-1-Ethyl]Carbamoyl N^2^-methyl-2-aminoethenoadenosine (7b)

MD-3-41 (250 mg, 0.69 mmol) and 4-nitrophenyl chloroformate (220 mg, 1.10mmol) were added to a round bottom flask and dissolved in pyridine (10 ml). The reaction was stirred for 2.5 hr at room temperature. To the reaction mixture was added tryptamine (221 mg, 1.38 mmol) dissolved in pyridine (2 ml). The reaction was stirred overnight at RT. The reaction was concentrated under reduced pressure. The crude mixture was taken up in ethyl acetate and sodium bicarbonate. The organic layer was extracted twice and combined. The organic layer was washed with sodium bicarbonate, brine and water. The organic layer was collected, dried over MgSO4, filtered, and dried under reduced pressure. The product was purified using flash chromatography (0-15% DCM/MeOH). The relevant fractions were pooled and concentrated under reduced pressure. Yield: 142 mg, 38%. The ^1^H NMR spectrum was (DMSO-d_6_): δ 1.36 (s, 3H), 1.57 (s, 3H), 2.80 (t, 2H), 3.07 (d, 3H), 3.22 (m, 4H), 4.08 (m, 2H), 4.25 (dd, 1H), 4.35 (td, 1H), 5.10 (dd, 1H), 5.54 (dd, 1H), 6.26 (d, 1H), 6.96 (t, 1H), 7.06 (m, 1H), 7.14 (d, 1H), 7.33 (d, 1H), 7.39 (t, 1H), 7.45 (d, 1H), 7.50 (d, 1H), 7.90 (q, 1H), 7.96 (d, 1H), 8.09 (s, 1H), 10.79 (m, 1H). ^13^C-DMSO-d_6_: δ 25.63, 25.89, 27.47, 28.72, 41.68, 49.07, 64.35, 81.99, 83.76, 84.48, 89.52, 109.61, 111.81, 112.02, 113.77, 116.67, 118.64, 118.67, 121.35, 123.09, 127.65, 132.23, 136.67, 137.24, 140.24, 142.17, 145.04, 156.18. ESI-MS [M+H] 547.4.

#### Synthesis of 2’,3’-Isopropylidine-5’-O-[(3-Indolyl)-1-Ethyl]Carbamoyl Ethenoadenosine (7c)

MD-3-41 (250 mg, 0.69 mmol) and 4-nitrophenyl chloroformate (307 mg, 0.96 mmol) were added to a round bottom flask and dissolved in pyridine (10 ml). The reaction was stirred for 2.5 hr at room temperature. To the reaction mixture was added tryptamine (310 mg, 1.94 mmol) dissolved in pyridine (4 ml). The reaction was stirred overnight at RT. The reaction was concentrated under reduced pressure. The crude mixture was taken up in ethyl acetate and sodium bicarbonate. The organic layer was extracted twice and combined. The organic layer was washed with sodium bicarbonate, brine and water. The organic layer was collected, dried over MgSO4, filtered, and dried under reduced pressure. The product was purified using flash chromatography (0-15% DCM/MeOH). The relevant fractions were pooled and concentrated under reduced pressure. The product was carried forward as mixture of starting material and product. Crude yield: 169 mg, 36%. ESI-MS [M+H] 517.3.

#### 5’-O-[(3-Indolyl)-1-Ethyl]Carbamoyl 2-aminoethenoadenosine (8)

MD-3-45 (74 mg, 0.14 mmol) was added to a scintillation vial with 4:1 TFA/H_2_O (2 ml). The reaction stirred at RT for 30 min. The reaction was transferred to a round bottomed flask with 0.1% TEA/Ethanol and subsequently concentrated under reduced pressure. 3 rounds of 50 ml of 0.1% TEA/Ethanol were used to quench the reaction. The crude product was purified using reverse phase flash chromatography 85-0% H_2_O/ACN). The relevant fractions were concentrated under reduced pressure to remove any acetonitrile, then flash frozen and lyophilized to yield the product as a white powder. Yield: 56.5 mg, 83%. The ^1^H NMR spectrum was (DMSO-d_6_): δ 2.76 (t, 2H), 3.20 (m, 2H), 4.05 (m, 3H), 4.19 (dd, 1H), 4.55 (t, 1H), 5.30 (m, 1H), 5.47 (s, 1H), 5.85 (d, 1H), 6.90 (t, 1H), 6.99 (t, 1H), 7.08 (d, 1H), 7.26 (d, 1H), 7.39 (t, 1H), 7.45 (d, 1H), 7.52 (s, 1H), 7.73 (d, 2H), 7.95 (t, 1H), 8.16 (s, 1H), 10.73 (s, 1H). ^13^C-DMSO-d_6_: ^13^C NMR (126 MHz, DMSO) δ 9.08, 26.00, 41.68, 46.21, 64.52, 71.12, 73.58, 82.94, 87.04, 110.60, 111.84, 112.05, 115.17, 118.65, 118.71, 121.38, 123.11, 127.65, 136.69, 137.12, 141.50, 145.59, 156.47. ESI-MS [M+H] 492.3.

#### Synthesis of 5’-O-[(3-Indolyl)-1-Ethyl]Carbamoyl N^2^-methyl-2-aminoethenoadenosine (9)

MD-3-47 was added to a scintillation vial with 4:1 TFA/H_2_O. The reaction stirred at RT for 30 min. The reaction was transferred to a round bottomed flask with 0.1% TEA/Ethanol and subsequently concentrated under reduced pressure. 3 rounds of 50 ml of 0.1% TEA/Ethanol were used to quench the reaction. The crude product was purified using reverse phase flash chromatography 85-0% H_2_O/ACN). The relevant fractions were concentrated under reduced pressure to remove any ACN, then flash frozen and lyophilized to yield the product as a white powder. Yield: 64.5 mg, 69%. The ^1^H NMR spectrum was (DMSO-d_6_): δ 2.75 (dd, 2H), 2.99 (d, 3H), 3.19 (m, 2H), 4.02 (dt, 1H), 4.06 (dd, 1H), 4.16 (q, 1H), 4.23 (dd, 1H), 4.68 (q, 1H), 5.32 (d, 1H), 5.43 (d, 1H), 5.87 (d, 1H), 6.90 (t, 1H), 6.99 (m, 1H), 7.07 (d, 1H), 7.25 (d, 1H), 7.37 (m, 2H), 7.45 (d, 1H), 7.76 (q, 1H), 7.88 (d, 1H), 8.05 (s, 1H), 10.72 (d, 1H). ^13^C-DMSO-d_6_: δ 25.98, 28.71, 41.68, 64.72, 71.26, 73.30, 82.73, 87.82, 109.52, 111.82, 112.06, 116.49, 118.65, 118.69, 121.36, 123.09, 127.66, 132.05, 136.68, 136.97, 141.19, 142.25, 145.00, 156.44. ESI-MS [M+H] 506.3.

#### 5’-O-[(3-Indolyl)-1-Ethyl]Carbamoyl Ethenoadenosine (10)

MD-3-45 (150 mg, 0.30 mmol) was added to a scintillation vial with 4:1 TFA/H_2_O (2.5 ml). The reaction stirred at RT for 30 min. The reaction was transferred to a round bottomed flask with 0.1% TEA/Ethanol and subsequently concentrated under reduced pressure. 3 rounds of 50 ml of 0.1% TEA/Ethanol were used to quench the reaction. The crude product was purified using reverse phase flash chromatography 85-0% H_2_O/ACN). The relevant fractions were concentrated under reduced pressure to remove any acetonitrile, then flash frozen and lyophilized to yield the product as a white powder. Yield: 96.5 mg, 69%. The ^1^H NMR spectrum was (DMSO-d_6_): δ 2.75 (t, 2H), 3.19 (m, 2H), 4.09 (m, 3H), 4.22 (dd, 1H), 4.61 (q, 1H), 5.36 (d, 1H), 5.53 (d, 1H), 6.00 (d, 1H), 6.90 (t, 1H), 6.98 (t, 1H), 7.08 (d, 1H), 7.26 (d, 1H), 7.39 (t, 1H), 7.45 (d, 1H), 7.50 (d, 1H), 8.02 (d, 1H), 8.47 (s, 1H), 9.24 (s, 1H), 10.73 (s, 1H). ^13^C-DMSO-d_6_: δ 25.99, 41.69, 71.07, 73.94, 83.15, 87.88, 111.83, 112.06, 112.73, 118.66, 118.71, 121.37, 123.10, 123.51, 127.66, 133.25, 136.69, 137.66, 139.05, 140.37, 140.94, 156.45. ESI-MS [M+H] 477.3.

#### Synthesis of 2’,3’-Isopropylidine-5’-O-(Sulfamoyl) 2-aminoethenoadenosine (11a)

MD-3-49 was added to a dry round bottom flask, purged with nitrogen gas, and cooled to 0°C. to the flask was added MD-3-1. TEA was added dropwise to the flask. The reaction was stirred at 0°C for 10 minutes, then RT for hour. The crude mixture was concentrated under reduced pressure and then purified by reverse phase chromatography (95-0% H_2_O). The relevant fractions were pooled and acetonitrile was removed under reduced pressure. The resulting solution was lyophilized producing the product as a white powder. Yield: 123 mg, 50%. The ^1^H NMR spectrum was (DMSO-d_6_): 1.29 (s, 3H), 1.49 (s, 3H), 2.47 (s, 3H), 4.06 (dd, 1H), 4.18 (dd, 1H), 4.31 (m, 1H), 5.22 (dd, 1H), 5.31 (dd, 1H), 6.17 (d, 1H), 7.37 (d, 1H), 7.49 (s, 2H), 7.55 (s, 2H), 7.87 (d, 1H), and 7.99 (s, 1H). ^13^C-DMSO-d_6_: 145.56, 142.30, 140.30, 136.83, 132.14, 116.70, 113.87, 110.14, 89.30, 84.68, 84.26, 81.74, 68.96, 40.90, 40.58, 40.49, 40.41, 40.32, 40.25, 40.16, 40.08, 39.99, 39.91, 39.82, 39.66, 39.49, 27.47, 25.77. ESI-MS [M+H] 440.3.

#### Synthesis of 2’,3’-Isopropylidine-5’-O-(Sulfamoyl) N^2^-methyl-2-aminoethenoadenosine (11b)

MD-3-49 (200 mg, 0.56 mmol) was added to a dry round bottom flask with DMF (2 ml), purged with nitrogen gas, and cooled to 0°C. to the flask was added MD-3-1 (259 mg, 2.24 mmol). TEA (78 µl, 0.56 mmol) was added dropwise to the flask. The reaction was stirred at 0°C for 10 minutes, then RT for 1 h. The crude mixture was concentrated under reduced pressure and then purified by reverse phase chromatography (95-0% H_2_O). The relevant fractions were pooled and acetonitrile was removed under reduced pressure. The resulting solution was lyophilized producing the product as a white powder. Yield: 91 mg, 38%. The ^1^H NMR spectrum was (DMSO-d_6_): 1.30 (s, 3H), 1.50 (s, 3H), 2.47 (s, 1H), 3.00 (d, 3H), 4.07 (dd, 1H), 4.17 (dd, 1H), 4.32 (m, 1H), 5.09 (dd, 1H), 5.49 (dd, 1H), 6.22 (d, 1H), 7.38 (d, 1H), 7.50 (s, 2H), 7.83 (q, 1H), 7.88 (d, 1H), and 8.01 (s, 1H). ^13^C-DMSO-d_6_: 145.08, 142.15, 140.11, 137.38, 132.23, 116.72, 113.98, 109.62, 89.59, 84.13, 83.81, 81.73, 68.51, 28.78, 27.39, and 25.59 ppm. ESI-MS [M+H] 426.3.

#### 2’,3’-Isopropylidine-5’-O-[N-(3-Indolepropionic acid)sulfamoyl] 2-aminoethenoadenosine Triethylammonium salt (12a)

MD-3-53 (120 mg, 0.37 mmol) and MD-3-17 (160 mg, 0.56 mmol) were added to a dry round bottom flask with DMF (2 ml). The flask was purged with nitrogen gas/vacuum and cooled to 0°C. To the flask was added DBU (70 µl, 0.47 mmol). The reaction stirred for 1 hour at 0°C and then overnight at room temperature. The crude mixture was concentrated under reduced pressure and then purified by reverse phase chromatography (95-0%, 1% TEA in H_2_O/ACN). The relevant fractions were pooled and concentrated under reduced pressure. Yield: 190 mg, 74%. The ^1^H NMR spectrum was (DMSO-d_6_): 1H NMR (500 MHz, DMSO) δ 1.31 (s, 3H), 1.53 (s, 3H), 2.38 (td, 2H), 2.85 (t, 2H), 3.90 (dd, 1H), 4.35 (td, 1H), 4.43 (dd, 1H), 5.21 (dd, 1H), 5.32 (dd, 1H), 6.14 (d, 1H), 6.87 (m, 1H), 7.00 (ddd, 1H), 7.04 (d, 1H), 7.28 (d, 1H), 7.42 (m, 2H), 7.79 (s, 2H), 7.93 (d, 1H), 8.11 (s, 1H), 9.47 (s, 0H), 10.66 (d, 1H). ^13^C-DMSO-d_6_: ^13^C NMR (126 MHz, DMSO) δ 9.42, 19.31, 21.98, 23.76, 25.68, 26.35, 27.56, 28.68, 32.17, 38.08, 46.22, 48.34, 53.85, 67.09, 82.20, 84.02, 84.50, 90.17, 110.07, 111.68, 113.39, 115.18, 116.60, 118.41, 118.64, 118.66, 121.15, 122.35, 127.63, 132.10, 136.67, 136.76, 136.91, 140.42, 142.36, 145.54, 165.86, 178.16. ESI-MS [M+H] 611.4.

#### Synthesis of 2’,3’-Isopropylidine-5’-O-[N-(3-Indolepropionic acid)sulfamoyl] N^2^-methyl-2-aminoethenoadenosine (12b)

MD-3-53 (160 mg, 0.36 mmol) and MD-3-17 (155 mg, 0.54 mmol) were added to a dry round bottom flask with DMF (2 ml). The flask was purged with nitrogen gas/vacuum and cooled to 0°C. To the flask was added DBU (59 µl, 0.40 mmol). The reaction stirred for 1 hour at 0°C and then overnight at room temperature. The crude mixture was concentrated under reduced pressure and then purified by reverse phase chromatography (95-0%, 1% TEA in H_2_O/ACN). The relevant fractions were pooled and concentrated under reduced pressure. Yield: 167 mg, 66%. The ^1^H NMR spectrum was (DMSO-d_6_): δ 1.35 (s, 3H), 1.57 (s, 3H), 2.34 (m, 2H), 2.83 (m, 2H), 3.07 (d, 3H), 4.02 (dd, 1H), 4.08 (dd, 1H), 4.37 (td, 1H), 5.12 (dd, 1H), 5.48 (dd, 1H), 6.22 (d, 1H), 6.93 (m, 1H), 7.04 (m, 2H), 7.30 (d, 1H), 7.45 (m, 2H), 7.87 (q, 1H), 7.95 (d, 1H), 8.14 (s, 1H), 10.68 (m, 1H). ^13^C-DMSO-d_6_: ^13^C NMR (126 MHz, DMSO) δ 12.25, 22.01, 28.78, 46.18, 67.73, 71.62, 73.48, 83.19, 87.80, 109.43, 111.66, 115.34, 116.46, 118.46, 118.72, 121.14, 122.36, 127.69, 132.08, 136.67, 137.04, 141.19, 142.33, 144.94, 177.92. ESI-MS [M+H] 597.4.

#### Synthesis of 5’-O-[N-(3-Indolepropionic acid)sulfamoyl] 2-aminoethenoadenosine Triethylammonium salt

**(13).** MD-3-69 (100 mg, 0.14 mmol) was added to a scintillation vial with 4:1 TFA/H_2_O. The reaction stirred at RT for 30 min. The reaction was transferred to a round bottomed flask with 0.1% TEA/Ethanol and subsequently concentrated under reduced pressure. 3 rounds of 50 ml of 0.1% TEA/Ethanol were used to quench the reaction. The crude product was purified using reverse phase flash chromatography 95-0% 0.1% in H_2_O/ACN). The relevant fractions were concentrated under reduced pressure to remove any ACN, then flash frozen and lyophilized to yield the product as a white powder. Yield: 95 mg, quant. The ^1^H NMR spectrum was (DMSO-d_6_): δ 2.40 (t, 2H), 2.51 (s, 3H), 2.87 (d, 2H), 4.02 (dd, 1H), 4.09 (s, 1H), 4.22 (s, 1H), 4.28 (dd, 1H), 4.69 (q, 1H), 5.29 (d, 1H), 5.43 (d, 1H), 5.90 (d, 1H), 6.93 (t, 1H), 7.03 (t, 1H), 7.08 (s, 1H), 7.31 (d, 1H), 7.43 (s, 1H), 7.48 (d, 1H), 7.63 (s, 2H), 7.94 (s, 1H), 8.15 (s, 1H), 10.68 (s, 1H). ^13^C NMR (126 MHz, DMSO): δ 9.31, 19.31, 21.96, 23.76, 26.35, 28.68, 32.17, 38.09, 46.21, 48.33, 53.85, 67.69, 71.48, 73.76, 83.26, 87.42, 109.99, 111.67, 115.25, 116.36, 118.46, 118.74, 121.16, 122.40, 127.67, 132.02, 136.46, 136.67, 141.39, 142.48, 145.41, 165.85, 177.92.

#### 5’-O-[N-(3-Indolepropionic acid)sulfamoyl] N^2^-methyl-2-aminoethenoadenosine (14)

MD-3-69 (100 mg, 0.14 mmol) was added to a scintillation vial with 4:1 TFA/H_2_O. The reaction stirred at RT for 30 min. The reaction was transferred to a round bottomed flask with 0.1% TEA/Ethanol and subsequently concentrated under reduced pressure. 3 rounds of 50 ml of 0.1% TEA/Ethanol were used to quench the reaction. The crude product was purified using reverse phase flash chromatography 95-0% 0.1% in H_2_O/ACN). The relevant fractions were concentrated under reduced pressure to remove any ACN, then flash frozen and lyophilized to yield the product as a white powder. Yield: 73 mg, 77%. The ^1^H NMR spectrum was (DMSO-d_6_): δ 2.29 (m, 1H), 2.30 (d, 1H), 2.78 (m, 2H), 3.97 (dd, 1H), 4.02 (td, 1H), 4.08 (dd, 1H), 4.19 (td, 1H), 4.70 (q, 1H), 5.29 (d, 1H), 5.35 (d, 1H), 5.85 (d, 1H), 6.86 (t, 1H), 6.96 (m, 1H), 7.00 (d, 1H), 7.23 (d, 1H), 7.35 (d, 1H), 7.41 (d, 1H), 7.71 (q, 1H), 7.86 (d, 1H), 8.07 (s, 1H), 8.80 (s, 1H), 10.60 (d, 1H). ^13^C-DMSO-d_6_: δ 12.25, 22.01, 28.78, 46.18, 67.73, 71.62, 73.48, 83.19, 87.80, 109.43, 111.66, 115.34, 116.46, 118.46, 118.72, 121.14, 122.36, 127.69, 132.08, 136.67, 137.04, 141.19, 142.33, 144.94, 177.92. ESI-MS [M+H] 556.3.

### Continuous Fluorescence Assay

Kinetics experiments were conducted on the Cary Eclipse UV spectrophotometer. All experiments were performed in HINT1 assy buffer (20 mM HEPES, 2 mM CaCl_2_, pH 7.4) at ambient temperature. HINT1 was exchanged into HINT1 assay buffer using 10 kD cutoff spin columns. HINT1 concentrations were determined via NanoDrop absorbance using an extinction coefficient determined from ExPASy ProtParam tool. Procedure was adapted from previous work.^24^ All assays were performed in a 600 µl tapered quartz cuvette. HINT1 and inhibitor were incubated for 30 sec followed by substrate addition. The cuvette was immediately placed in the fluorimeter following substrate addition. The rate of fluorescence was measured over 5 minutes and the slope was determined. Fluorescence was converted to [Tryptamine] using the slope gathered from a tryptamine standard curve. The inhibition curves were plotted in Graphpad Prism and K_i_ values were determined using a nonlinear regression. Values are reported with the standard error of the residual.

### Protein Crystallography

Crystals were grown via hanging drop vapor diffusion, with drops composed of 2 μL of protein and 2 μL of well solution. Well solutions contained 10-14 % (w/v) PEG 4000 and 100 mM sodium cacodylate pH 6.0-6.6. Crystals were formed after 4 days of incubation at 8 °C. Co-crystals with inhibitors were prepared by soaking preformed crystals in mother liquor containing 12.5 mM of each ligand for 3-40 min. DMSO was used to adequately dissolve the inhibitors, which came at a cost to soaking crystal’s structural integrity and thus the ability to soak the crystals for long. Soaked crystals did not need cryopreservation, but had to be mounted on a very thin film from the crystallization buffer. The excess liquid was removed by gentle touching of the mounting loop against the plate surface and the crystal was then flash cooled directly in an N_2_ stream. Diffraction data were collected using Rigaku XtaLAB Synergy-S diffractometer equipped with a HyPix-6000HE Hybrid Photon Counting (HPC) detector and Cu microfocus sealed X-ray source as well as a low-temperature Oxford Cryostream 800 liquid nitrogen cooling system at 100 K. The data collection strategy was calculated within *CrysAlis PRO* to ensure desired data redundancy and percent completeness. Data were processed, integrated and scaled using *CrysAlis PRO* and AIMLESS Data acquisition and processing statistics are shown in Table X1.^25^

Molecular replacement was conducted with hHINT1 coordinates (PDB ID: 6yqm [X3]) using *MOLREP* software.^26^ Modeling and molecular visualization were performed in *COOT*.^27^ Ligand restraints were calculated using *JLigand* [X6] and refinement was performed using *REFMAC5*.^28^ All refinement steps were monitored with *R*_cryst_ and *R*_free_ values. The stereochemical quality of the resulting models was assessed using the program *MOLPROBITY* and the validation tools implemented in *COOT*.^29^ The values of the mean temperature factors for protein main and side chains, ligands and water molecules were calculated using the program BAVERAGE from CCP4 suite.^30^ Superposition of protein structures was performed using the program LSQKAB.^31^ The refinement statistics of the described structures are listed in **Table S1**.

### Animals

ICR-CD-1 mice (male and female, 21-30 grams) were maintained on a 12-hour light/dark cycle with unrestricted access to water and food. All experiments were approved by the Institutional Animal Care and Use Committee of the University of Minnesota. Animal experiments were adapted from our previous work.^4^

### Drug Preparation for Behavioral Assays

Morphine sulfate (NIDA) and endomorphin-2 (endo-2) were dissolved in 0.9% saline. Endomorphin-2 was prepared as previously described.^32^ Briefly, endo-2 was synthesized using solid-state methods and HPLC-purified by the Microchemical Facility of the University of Minnesota and was dissolved for intrathecal injection in 0.9% normal saline. All stocks of TrpGc and TrpGc analogues were dissolved in 5% DMSO, 10% EtOH, and 10% cremophor and diluted with diH20 to a final concentration of 0.5% DMSO, 1% cremophor, and 1% ethanol. From this stock, a final concentration was reached by diluting the stock solution with 0.9% normal saline into the injection concentration.

### Intrathecal Injections

All drugs were delivered in 5 μL volumes via intrathecal injection in conscious mice.^33^ Briefly, the mice were held by the iliac crest and drugs were injected into the intrathecal space by a 30-gauge, 0.5-inch needle attached to a 50 μL Luer-hub Hamilton syringe.

### Warm Water Tail Immersion Assay

Antinociception was measured using a warm water tail immersion assay. Mice were wrapped in a soft cloth with their tails exposed and approximately 3/4 of the tail was dipped into a warm water bath (49 or 52.5°C). The latency for the mouse to flick its tail was recorded before and after intrathecal administration of drug. In order to avoid tissue damage, a maximum cutoff of 12 seconds was set. A minimum of 4 mice were used for each drug, and each subject received only one HINT1 inhibitor compound.

### Morphine Inhibition of NMDA-evoked Behavior

Intrathecally-injected NMDA gives rise to both a transient thermal hypersensitivity that can be measured by a warm water tail immersion assay, and a caudally directed scratching and biting behavior lasting for 1-5 minutes. For this initial screen of the TrpGc analog compounds, we measured the impact of each inhibitor on morphine’s inhibition of NMDA-evoked transient thermal hypersensitivity. A baseline tail flick latency (pre-NMDA tail flick latency) was recorded at 49°C. TrpGc or a TrpGc analog was intrathecally injected (0.1-30 nmol/5 μL) into male ICR mice (25-30g) 10 minutes prior to an intrathecal injection of morphine sulfate (10 nmol/5 μL). After a period of 10 minutes, NMDA was injected intrathecally and another tail flick latency was recorded (post-NMDA tail flick latency). The percent maximum possible effect (%MPE) was calculated according to the following equation:

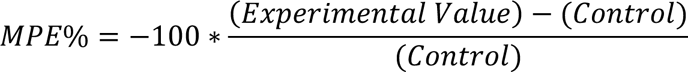

where the Control value is the average reduction of tail flick latency within the cohort of subjects receiving only NMDA treatment:

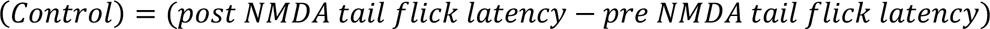

and the Experimental Value is the change in tail flick latency in the presence of NMDA + morphine or NMDA + morphine + HINT1 inhibitor.

### Endomorphin-2 tolerance

Baseline thermal responsiveness was assessed in a 52.5°C water bath tail-immersion assay with a cutoff time of 12 seconds. TrpGc, TrpGc analog or vehicle was injected intrathecally, followed 5 minutes later by an intrathecal injection of endomorphin-2 (endo-2) at a dose of 10 nmol/5 μL into male and female ICR mice. Observation of a Straub tail for each subject was used as an indication of a successful intrathecal injection of an opioid agonist. Thirty minutes following this injection, an additional tail flick was assessed in order to confirm a return to baseline responsiveness and a lack of continued analgesia (predrug latency). A probe dose of endo-2 (10 nmol/5 μL, i.t.) was injected, and a final tail flick latency (postdrug latency) was assessed 2.5 minutes following this probe endo-2 injection.

The results are expressed as a percentage maximum possible effect (%MPE) according to the following equation:

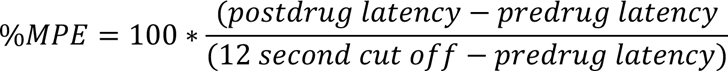

### Motor Coordination

Motor coordination was assessed by rotarod assay, as previously described.^33^ Briefly, after a training session, mice were injected intrathecally with either vehicle control or a compound of interest and placed upon a rotating rod (Ugo Basile, Carese, Italy) 5 minutes following this injection. The rod accelerated from 4-40 rpm over 300 seconds. The latency to fall was recorded and compared between the subjects that received vehicle control and subjects that received a compound of interest. Data are presented as mean +/- SEM.

### Open Field Activity Monitoring

Sedation and motor activity were assessed by the open field assay (ENV-510S-A from MedAssociates, Fairfax, VT, USA). Mice were intrathecally injected with either vehicle control or a compound of interest and placed in an open field chamber 5 minutes following injection. Subjects’ movement was recorded for 10 minutes and compared between the vehicle control and compound of interest for the total distance traveled, the average speed, and the time spent at rest. Data are presented as mean +/- SEM.

### Behavioral Data Analysis

For behavioral assays, data were calculated as described above. The data are represented as mean +/- SEM for each assay. A minimum of three doses were used for dose-response analysis.

#### Effect of HINT1 inhibition on HINT1 and p53 expression

BT549 cells were obtained from ATCC cultured in RPMI 1640 (Gibco, 11875093) and SY-SY5Y cells, obtained from ATCC, were cultured in Dulbecco’s Modified Eagle Medium (DMEM, 11966025) at 37 °C in a humidified incubator under 5% CO_2_. Medium for both cell lines was supplemented with 10% fetal bovine serum (GenClone, 25-514) and GlutaMAX (ThermoFischer, 35050061). For BT549, RPMI 1640 was also supplemented with 5 μg/mL insulin (Sigma-Aldrich, I9278). BT549 and SH-SY5Y were grown in two separate T75 flasks for each cell line. BT549 were grown to ∼4 million/mL in 10 mL and SH-SY5Y were grown to ∼10 million/mL in 10 mL medium. Cells were then plated at ∼ 1 million cells/well into two separate 6-well plates and allowed to attach overnight. TrpGC stock was prepared in DMSO (Sigma Aldrich, D8418) for a final concentration of 5mM TrpGC in DMSO. Working stock solution of DMSO in RPMI 1640 and DMEM with was then prepared for a final DMSO concentration of 0.2% DMSO. 100 µM TrpGC with a final DMSO concentration of 0.2% was prepared separately using both medias. 0.2% DMSO working stock was then used to dilute and obtain 50, 25 and 12.5 µM TrpGC. Wells were designated as media, DMSO control (0.2%), 100, 50, 25 and 12.5 µM TrpGC. 5mL of each stock was added in the designated wells and cells were incubated with the indicated treatments for 72 hours. After 72 hours, cells were collected through trypsinization (Gibco, 25200072). Media was discarded and ice-cold RIPA buffer containing Pierce™ EDTA-free protease inhibitor tablet (ThermoFischer, A32965) (1 tablet/10mL) was used to resuspend the cell pellets (∼100 µL/pellet). After 5-10 minutes of incubating pellets in ice with RIPA, contents were sonicated for 30 minutes, followed by high-speed centrifugation for 10 minutes. Lysate concentrations were determined using Pierce™ BCA Protein Assay Kit (ThermoFischer, 23227). Cell lysates from each sample were prepared and electrophoresed on a 10-well 4–12% NuPAGE gradient gel (Invitrogen, NP0321BOX) and transferred onto 0.45 μm pore polyvinylidene difluoride membranes (PVDF) (Invitrogen, LC2006). Immunoblotting was performed with anti-HINT1 (Bethyl, A305-373A, 1:2000, rabbit), anti-p53 (Invitrogen, MA5-14067, 1:1000, mouse) and anti-β-actin (Sigma-Aldrich, A1978, 1:2000, mouse) primary antibodies. The membrane was incubated with primary antibodies overnight at 5 °C followed by goat anti-rabbit IgG (AlexaFluor 546) conjugated secondary antibody (Invitrogen, A11010, 1:1500) and goat-anti-mouse AlexaFluor 680 Plus (Invitrogen, A32729, 1:1500). Channels 600 and 700 were used to image the membranes using Odyssey Fc Imaging system (Li-Cor). Band intensity was quantified using ImageJ and GraphPad Prism software was used to obtain mean ± SEM from three biological replicates.

## Acknowledgements

Financial support from the University of Minnesota Foundation is gratefully acknowledged.

