## Supplemental Data for "HINT1 Inhibitors as Selective Modulators of MOR-NMDAR Cross Regulation and Non-Opioid Analgesia"

Centre of Molecular and Macromolecular Studies[†],
Polish Academy of Sciences
Sienkiewicza 112, 90-363 Lodz, Poland

---

**HINT1 Inhibition Curves**

**Figure S1**

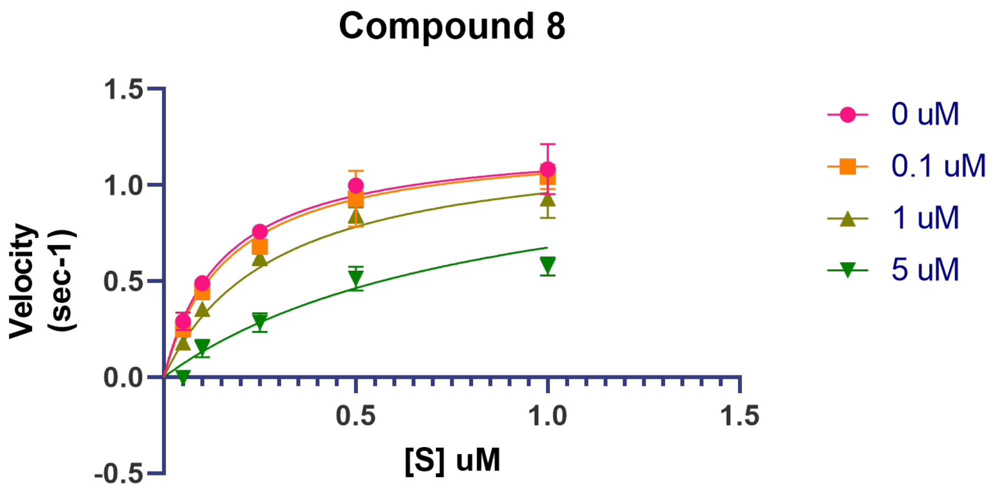

**Figure S2**

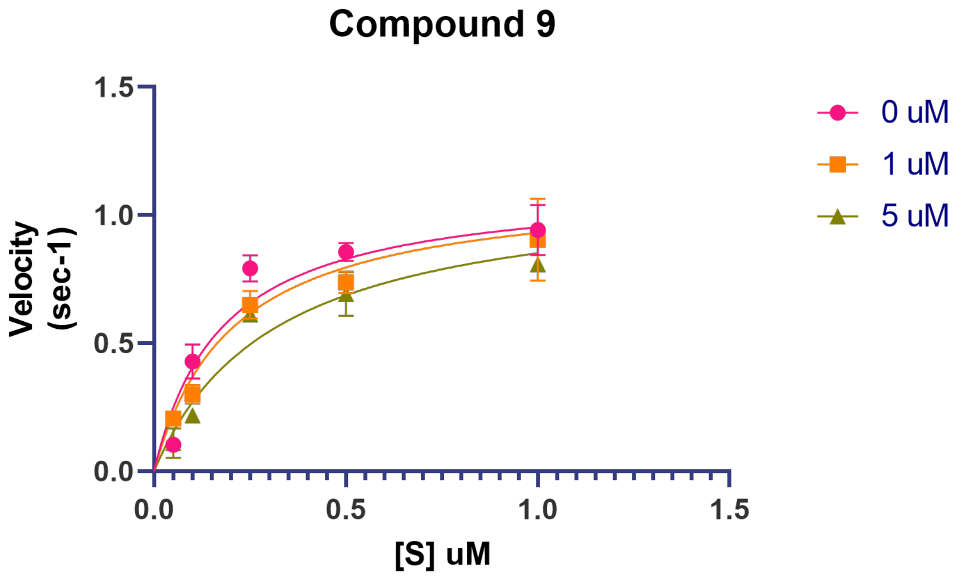

**Figure S3**

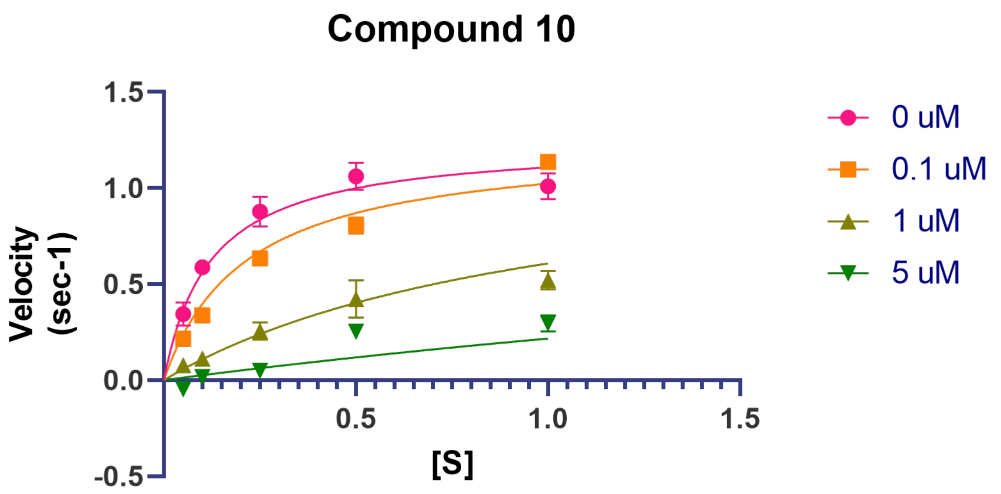

**Figure S4**

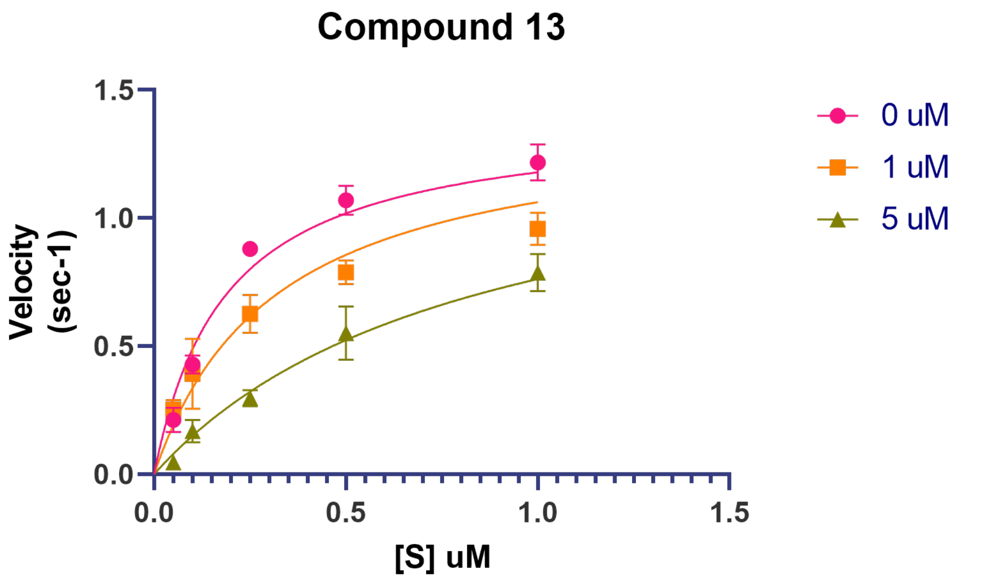

**Figure S5**

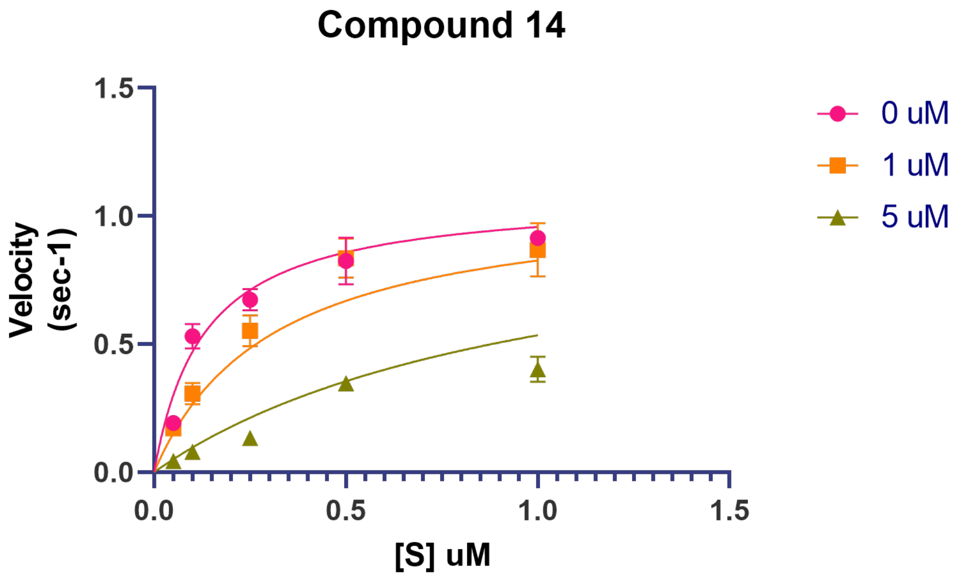

**Figure S6**

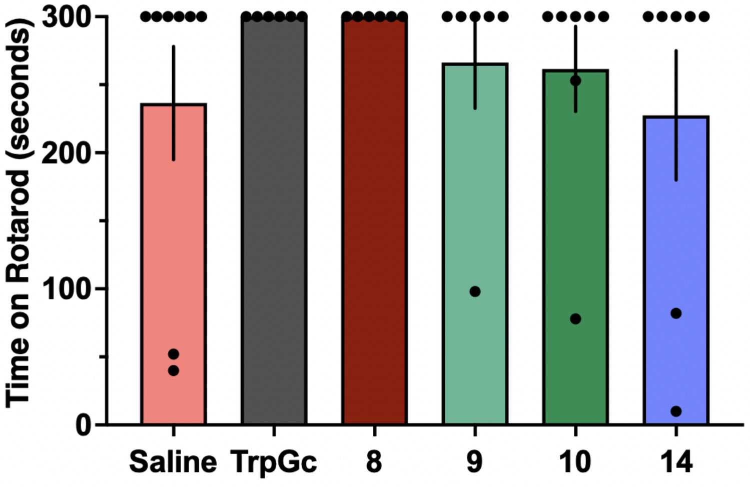

**Figure S6: The inhibitors do not induce motor coordination impairment as measured by the rotarod assay.** All compounds were administered intrathecally at a dose of 10 nmol 5 minutes prior to placement on the rotating rod, and the latency to fall off the rod was recorded in seconds.

There was no difference between treatment groups as compared to the saline control. Ordinary one-way ANOVA followed by Dunnett's multiple comparisons test. N = 6-8/group, m + f.

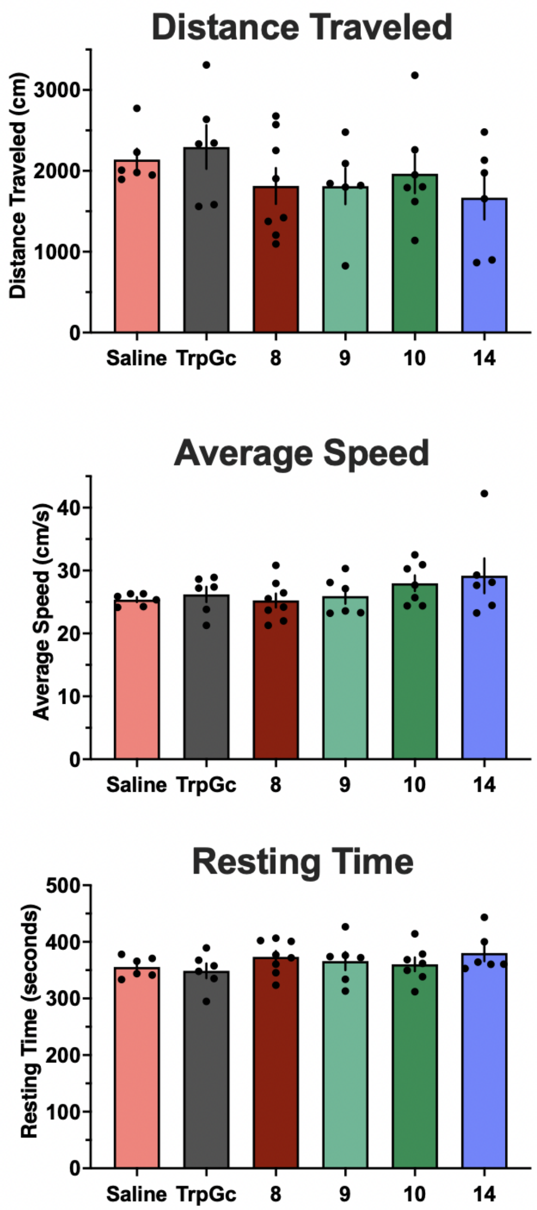

**Figure S7: The inhibitors do not induce alterations in locomotion as measured by the open field assay.** All compounds were administered intrathecally at a dose of 10 nmol 5 minutes prior to placement in an open field apparatus. Total movement was monitored for 10 minutes. The distance traveled (top panel, cm), average speed (middle panel, cm/s), and resting time (bottom panel, seconds) were recorded and compared to a cohort of mice that received an injection of intrathecal saline as a control. No differences for any parameter were observed for any compound. Ordinary one-way ANOVA followed by Dunnett's multiple comparisons test. N = 6-8/group, m + f.

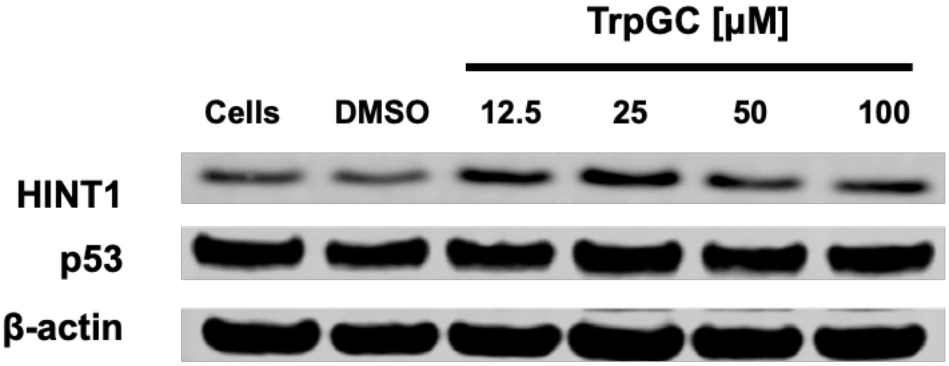

**Figure S8.** Representative western blot image showing the effect of HINT1 inhibition on p53 expression. BT549 cells were incubated for 72 hours with up to 100 µM of variable concentrations of inhibitor, TrpGC. No significant difference in HINT1 or p53 expression was observed.

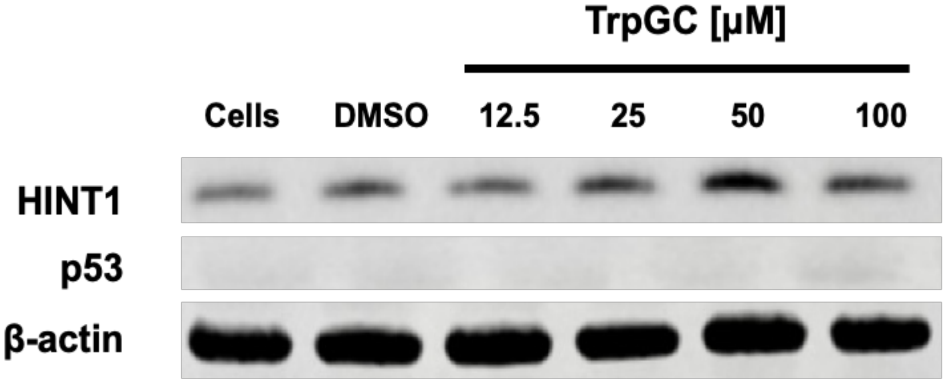

**Figure S9.** Representative western blot image showing the effect of HINT1 inhibition in SH-SY5Y cells. No p53 expression was observed over 72 hours of incubation. No significant difference in HINT1 expression was observed.

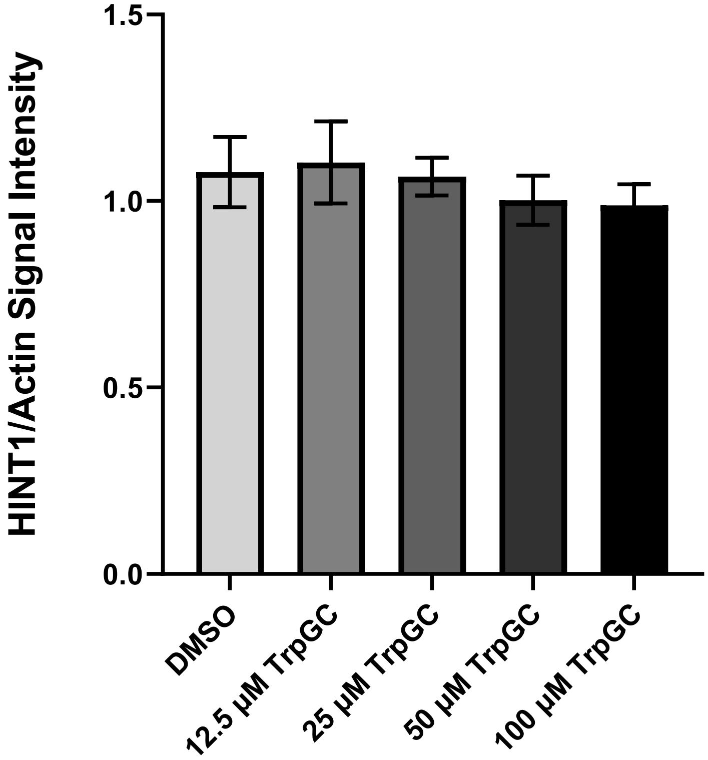

**Figure S10**. Densitometry analysis was performed using ImageJ and HINT1 band density ratio to β-actin control were determined. Statistical analysis was performed using Brown-Forsythe and Welch ANOVA followed by Dunnett's T3 multiple comparisons test. Data shown as mean ± SEM (n=3 individual experiments). No significant difference among treatment means was determined ($p>0.05$).

**Table S1. Crystallographic parameters and data collection statistics.**

| PDB ID | 8P8P | 8PA6 | 8PA9 | 8PAF | 8PAI |
|---|---|---|---|---|---|
| Crystallization conditions | 10 % w/v PEG4000, 0.1 M sodium cacodylate pH 6.0 | 12 % w/v PEG4000, 0.1 M sodium cacodylate pH 6.0 | 12 % w/v PEG4000, 0.1 M sodium cacodylate pH 6.0 | 14 % w/v PEG4000, 0.1 M sodium cacodylate pH 6.0 | 10 % w/v PEG4000, 0.1 M sodium cacodylate pH 6.0 |
| Crystal size (µm) | 70 × 40 × 40 | 70 × 40 × 40 | 70 × 40 × 40 | 70 × 40 × 40 | 70 × 40 × 40 |
| Ligand | TrpEtAdC | Trp2AEtAdC | Trp2MAEtAdC | Trp2AEtAdAS | Trp2MAEtAdAS |
| Ligand code | X7I | XKB | XKF | XKK | XKO |
| Soaking time (min.) | 10 | 10 | 20 | 3 | 5 |
| X-ray source | Rigaku XtaLAB Synergy-S | | | | |
| Wavelength (Å) | 1.54184 | | | | |
| Detector | HyPix-6000HE | | | | |
| Detector distance (mm) | | | | | |
| Oscillation width (°) | 0.38 | 0.38 | 0.38 | 0.38 | 0.38 |
| Temperature (K) | 100 | 100 | 100 | 100 | 100 |
| No. of frames | 609 | 1326 | 1316 | 788 | 989 |
| Space group | *C*2 | *C*2 | *C*2 | *C*2 | *C*2 |
| Unit-cell parameters | | | | | |
| *a* (Å) | 78.73 | 78.84 | 78.60 | 79.19 | 77.67 |
| *a* (Å) | 46.55 | 46.32 | 46.31 | 46.55 | 46.43 |
| *c* (Å) | 64.10 | 63.97 | 64.01 | 64.11 | 63.79 |
| *α* (°) | 90.00 | 90.00 | 90.00 | 90.00 | 90.00 |
| *β* (°) | 94.74 | 94.80 | 94.71 | 94.84 | 94.62 |
| *γ* (°) | 90.00 | 90.00 | 90.00 | 90.00 | 90.00 |
| Total no. of reflections | 71108 (3434) | 169719 (5651) | 186288 (5964) | 72260 (6027) | 119678 (4221) |
| Unique reflections | 18317 (1200) | 31671 (1554) | 36820 (1815) | 13742 (1116) | 21127 (1216) |

| | | | | | |
|---|---|---|---|---|---|
| Completeness (%) | 99.6 (98.7) | 99.9 (100) | 99.8 (98.9) | 99.8 (99.9) | 99.8 (99.8) |
| Resolution (Å) | 19.22-1.90 (1.94-1.90) | 19.10-1.58 (1.61-1.58) | 18.45-1.50 (1.53-1.50) | 18.48-2.10 (2.16-2.10) | 19.04-1.80 (1.84-1.80) |
| $R_{merge}$[a] | 0.100 (0.331) | 0.074 (0.661) | 0.062 (0.444) | 0.082 (0.229) | 0.087 (0.369) |
| $R_{p.i.m}$ | 0.057 (0.220) | 0.035 (0.396) | 0.028 (0.315) | 0.039 (0.108) | 0.039 (0.226) |
| Multiplicity | 3.9 (2.9) | 5.4 (3.6) | 5.1 (3.3) | 5.3 (5.4) | 5.7 (3.5) |
| Mosaicity | 1.34 | 1.19 | 1.32 | 1.53 | 1.47 |
| Wilson *B* factor | 9.2 | 8.2 | 9.0 | 10.3 | 10.0 |
| Mean *I*/sd(*I*) | 8.9 (2.5) | 15.3 (2.1) | 14.2 (2.3) | 14.9 (6.6) | 12.6 (3.0) |
| *CC(1/2)* | 0.994 (0.891) | 0.998 (0.742) | 0.999 (0.840) | 0.998 (0.963) | 0.997 (0.893) |

**Table S2. Refinement statistics.**

| PDB ID | 8P8P | 8PA6 | 8PA9 | 8PAF | 8PAI |
|---|---|---|---|---|---|
| No. of reflections used in refinement | 17388 | 30046 | 34937 | 13141 | 20130 |
| No. of reflections used to $R_{free}$ | 910 | 1625 | 1878 | 592 | 982 |
| $R_{cryst}$ ($R_{free}$) | 0.154 (0.190) | 0.147 (0.191) | 0.149 (0.179) | 0.136 (0.187) | 0.149 (0.191) |
| No. of non H-atoms | | | | | |
| Protein | 1887 | 1886 | 2000 | 1857 | 1901 |
| Solvent | 267 | 390 | 332 | 291 | 238 |
| Ligand | 35 | 36 | 37 | 39 | 40 |
| R.m.s.d.s from ideal values | | | | | |
| Bond lengths (Å) | 0.008 | 0.010 | 0.011 | 0.008 | 0.008 |
| Bond angles (°) | 1.461 | 1.540 | 1.657 | 1.461 | 1.528 |
| Ramachandran plot | | | | | |
| Favoured [%] | 98.7 | 98.7 | 99.1 | 99.1 | 98.7 |
| Allowed [%] | 1.3 | 1.3 | 0.9 | 0.9 | 1.3 |
| Outliers [%] | 0 | 0 | 0 | 0 | 0 |
